# RBP47 family members are negative regulators of heat stress tolerance in *Arabidopsis thaliana*

**DOI:** 10.1101/2025.05.18.654719

**Authors:** Itzell E. Hernández-Sánchez, Ibrahim Tarbiyyah, Alyssa Kearly, Federico Martinez-Seidel, Israel Maruri-Lopez, Yue Chen, Olivia S. Rissland, Wei Wang, Salim Sioud, Adrian Schwarzenberg, Anna C. Nelson Dittrich, Andrew DL Nelson, Monika Chodasiewicz

## Abstract

Stress granules (SGs) are liquid-liquid phase-separated condensates that sequester RNA, proteins, and metabolites to modulate cellular physiology under stress. In Arabidopsis, the RNA-binding protein RBP47b is a canonical SG marker, yet its functional contribution to thermotolerance remains unresolved. Here, we combined mTurboID proximity labeling with multi-omics profiling to define the RBP47b interactome and its physiological impact. mTurboID identified a stress-specific enrichment of 40S ribosomal subunits within RBP47b SGs, implicating these condensates in translational control. Surprisingly, quadruple mutant plants lacking all four RBP47 paralogues (*rbp47abcc′*) displayed enhanced survival, attenuated growth delay, and faster recovery of photosynthetic efficiency after severe heat stress. Integrated transcriptome, proteome, and metabolome analyses revealed that this gain of thermotolerance is associated with (i) accelerated re-initiation of translation, (ii) constitutively elevated jasmonate and oxylipin pools, and (iii) reduced ROS accumulation during heat and recovery. We conclude that the RBP47 family acts as a negative regulator of heat tolerance by sequestering 40S subunits and limiting translational restart; loss of these SG scaffolds pre-primes jasmonate-dependent detoxification pathways and expedites proteome rebuilding, thereby conferring superior thermotolerance.

## Introduction

Elevated temperatures represent a significant detrimental abiotic stress for plants, posing critical challenges to crop production and food security ^1,2^. Plants have evolved adaptive mechanisms to withstand such adverse conditions and respond effectively to negative stimuli. One notable mechanism is the formation of biomolecular condensates driven by liquid-liquid phase separation (LLPS) ^3,4^. Over the past decade, these condensates have received increasing attention for their roles in temperature responses and stress tolerance in plants ^5^. Stress granules (SGs) are transient biomolecular condensates formed in response to diverse biotic and abiotic stressors that dissolve once the stress subsides ^6^. SGs represent a conserved stress response mechanism across living organisms ^7^. However, the precise role of SGs is still unknown.

Emerging evidence indicates that SGs serve as pivotal regulatory hubs that rapidly sequester specific cytoplasmic mRNAs and proteins, allowing a faster re-wiring of the molecular machinery during stress. Once favorable conditions are restored, these sequestered molecules are either released, allowing mRNAs to resume translation and proteins to resume their functional roles, or targeted for degradation to optimize cellular homeostasis during recovery from heat stress ^8,9^.

In plants, several RNA-binding proteins (RBPs) have been identified as SG markers ^10–12^, including the RNA-binding protein 47b (RBP47b) ^13^. Recent findings indicate that RBP47b participates in SG formation in response to heat stress and phenolic acid (PA) treatment ^14^, a signaling molecule involved in plant-to-plant communication. RBP47b belongs to a four-member gene family consisting of RBP47 a, b, c, and c’, sharing high protein sequence similarity (>75%). Although the RBP47 family has been systematically reported as part of the SG interactome ^4^, the precise functions and regulatory mechanisms of RBP47 in thermotolerance remain poorly understood.

In the present study, we set out to define the molecular contribution of RBP47 proteins to thermotolerance. First, using proximity-dependent biotin labeling (TurboID), we mapped the RBP47b interactome in control, heat, and recovery conditions. RBP47b SGs selectively engage proteins involved in translation, photosynthesis, hormone (SA/JA) signaling, and detoxification. Next, focusing on Col-0 and *rbp47abcc′* quadruple mutants, the transcriptome, proteome, metabolome, and lipidome were analyzed in response to high-temperature stress.

The combined analyses reveal that RBP47 family members act as negative regulators of heat tolerance.

## Results

### Heat stress modulates RBP47b dynamics, leading to SG formation in Arabidopsis cells

To increase our understanding of RBP47b function in plants, we explored its cellular dynamics in the context of heat stress response. RBP47b was previously reported to localize to SGs under heat stress conditions ^13^. To study the behavior of RBP47b under control, heat, and recovery phases, Arabidopsis plants (Col-0) expressing *UBQ10p:RBP47b:eYFP* were generated. Gene expression levels were determined to be ∼6-fold above endogenous *RBP47b* gene in Col-0, and no phenotypic differences were detected relative to control plants **(Supplementary Figure 1A-C)**. Consistent with prior findings, RBP47b localized in the cytosol and nucleus under control conditions (23°C). Upon heat stress (42°C), the cytosolic RBP47b fraction rapidly re-localized into SGs within approximately five minutes, while its nuclear localization remained unchanged (**Figure 1A**). The number of SGs increased through the first 10-45 min following stress (**Figure 1B**). Upon removal of the heat stress, RBP47b dissociated from SGs after 240–300 minutes (**Figures 1C-D**). The formation of SGs was prevented by incubating with cycloheximide (CHX), a translation elongation inhibitor, prior to heat stress exposure, suggesting that the release of mRNA from polysomes is an essential mechanism for RBP47b-SGs assembly **(Figure 1A)**. These findings indicate that RBP47b follows the canonical behavior of other SG-associated proteins, rapidly assembling into SGs upon stress induction and gradually disassembling after stress removal.

**Figure 1.**
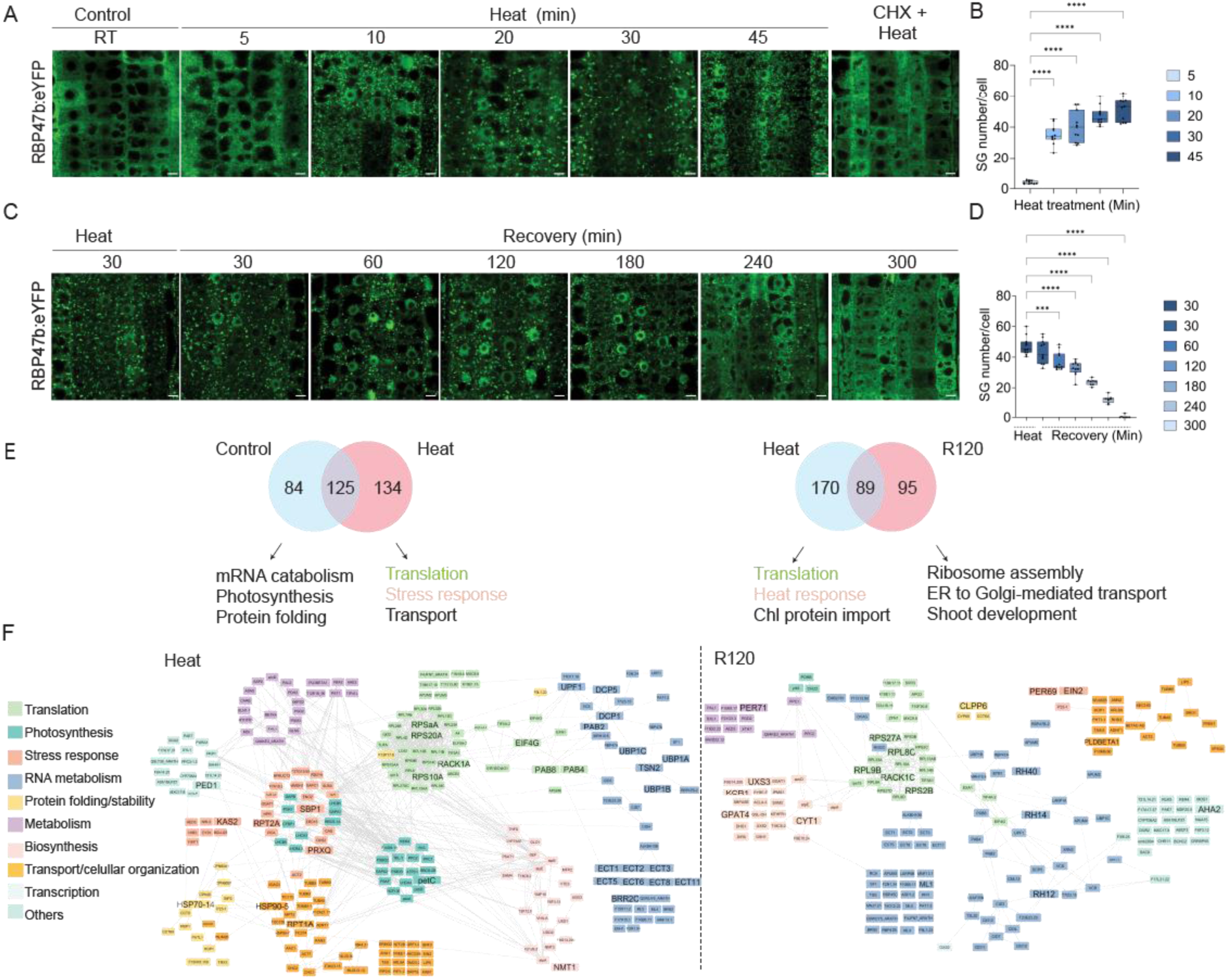
RBP47b is associated with SGs in response to high-temperature stress. **A.** SG dynamic of RBP47b-eYFP in 5-day-old Arabidopsis seedlings under control (23°C), heat (42°C) stress conditions, and CHX treatment/heat stress combination. **B.** Number of SGs per cell during the assembly phase. **C.** Disassembly dynamics of SG in the recovery phase after heat treatment. **D**. Number of SGs per cell during the disassembly phase. **E.** Venn diagram showing the overlap of proteins identified as potential interactions between control/heat (left) and heat/R60 (right). **F.** Interactome networks for heat and recovery conditions built in Cytoscape ^15^ based on STRING analysis with high-confidence interaction scores (0.7 cutoffs), including all types of evidence (see Materials and Methods). Data are presented as the mean ± SD of 11 biological replicates. Significant differences between time points are indicated by asterisks (*P ≤* 0.05, one-way ANOVA with post hoc Dunnett’s multiple test comparison).

### The high-resolution proxitome of RBP47b reveals dynamic protein networks under control, heat, and recovery phases

While the composition of RBP47b heat-induced SG cores in Arabidopsis has been characterized ^13^, a comprehensive view of how RBP47b’s interactions change during control, heat stress, and recovery conditions remains poorly understood. To bridge this gap, we adapted a previously established proximity labeling (PL) protocol for plants ^16^, refining it to address our specific biological question. We generated stable lines expressing UBQ10p:RBP47b–miniTurbo–mVenus (RBP47b^mTb^, line L6) or the ligase control UBQ10p:miniTurbo–NES–mVenus (NES^mTb^, line L5) in a Col-0 background; both transgenes accumulated 2- to 6-fold above endogenous *RBP47b* transcript levels **(Supplementary Figure 2A-B)**. Transgenic lines and Col-0 plants were phenotypically similar **(Supplementary Figure 2C)**. We then established 40 °C for 1h for heat treatment as the minimal heat pulse that re-localizes RBP47b^mTb^ to stress granules while leaving NES^mTb^ diffuse in the cytoplasm **(Supplementary Fig. 3A–B)**. Exogenous biotin application (30–100 µM) neither induced SGs at 22 °C nor inhibited their formation at 40 °C. Streptavidin blots revealed negligible background in Col-0 but robust labeling in RBP47b^mTb^ under 30 µM biotin. This concentration was adopted for all PL experiments **(Supplementary Fig. 3C–D)**. For the recovery time point, we selected 120 min (R120) based on our confocal RBP47b^mTb^ SG disassembly dynamics and biotinylation patterns **(Supplementary Figure 3E–F)** (For detailed description of proxitome optimization, refer to the Material and Methods section).

With the experimental conditions for the PL assay established, we focused on investigating the RBP47b interactome under control, heat stress, and recovery conditions (R120). A total of 4,457 proteins across all samples/treatments were identified by label-free quantitative proteomics **(Supplementary Table 1)**. After stringent data filtering, including the removal of background proteins present in NES^mTb^ and Col-0 (treated with 30 µM biotin) under control or heat conditions (see Materials and Methods), 209 proteins were found to be in close proximity to RBP47b under control conditions, 259 under heat, and 184 under recovery **(Supplementary Table 1)**. From these, 125 proteins are shared between control and heat, and 89 proteins are shared between heat and recovery **(Figure 1E)**. To understand the dynamics of RBP47b-associated proteins, interactome networks were generated for all conditions **(Figure 1F and Supplementary Figure 4)**. Gene Ontology (GO) enrichment analysis revealed that proteins identified across all subsets are associated with various biological processes, including RNA metabolism, stress response, photosynthesis, and lipid metabolism **(Supplementary Figure 5)**. Interestingly, shared protein components involved in SG assembly/disassembly and P-body assembly were identified during the transition from control to heat and heat to recovery. These findings suggest that RBP47b is consistently associated with proteins that regulate cytosolic condensate dynamics, reinforcing the model that RBP47b itself is a key regulator of SGs. Notably, among the shared protein components between control and heat stress conditions, we identified a distinct cluster involved in detoxification pathways and regulation of lipid metabolism, including CAT3, GGAT1, ABCG36, CYP73A5, METK4, accD, and F21J9.2 **(Supplementary Table 1)**. This highlights a potential link between RBP47b-associated SGs and the coordination of metabolic and stress-responsive processes during exposure to heat stress.

Focusing on proteins uniquely associated with each condition, we reported 84 proteins specific for control, 134 proteins specific for heat stress, and 95 proteins particularly related to the recovery phase **(Supplementary Table 1)**. Under control conditions, RBP47b is associated with stress response proteins (HSP70-1, HSP70-3, HSP90-6, DJ1A, RBP45b, NCL, and GSTF2), mRNA catabolic process (TSN1, XRN4, LARP1A, RH14, and MRG7.19), and cellular organization and transport (TUBB4 and TUBB7) **(Supplementary Figure 4 and Supplementary Table 1)**. Interestingly, many of these proteins have been reported as part of the pre-network SG interactions, interactions that exist before the stress^13,17–20^.

From the 134 potential RBP47b-associated proteins specific to heat conditions, we identified well-known SGs components, including TSN2, PAB2, PAB4, PAB8, UBP1A-C, and elongation factors such as EIF4G, as well as proteins associated with processing bodies (PBs) like DCP1 and DCP5 ^10,11,13,21,22^ **(Figure 1F and supplementary Table 1)**. These known SG constituents validated our approach, confirming its ability to capture SG-associated proteins. In addition to these proteins, a subset of proteins related to ROS-detoxification, hormonal pathways, and lipid metabolism were observed to be recruited explicitly under heat stress (NMT1, PRXQ, petC, SBP1, PED1, and KAS2) ^23–28^.

Interestingly, in addition to the previously reported m^6^A reader EVOLUTIONARILY CONSERVED C-TERMINAL REGION 2 (ECT2) ^13^, two other ECT family members, ECT1 and ECT8, were found associated explicitly with RBP47b under heat. Other members of the ECT family, ECT3, ECT5, ECT6, and ECT11, were found in proximity to RBP47b under both control and heat conditions. Under heat, beyond known SG components, we also identified proteins involved in stress response, RNA metabolism (BRRC2, UPF1), transport, protein folding (HSP70-14, HSP90-5), and protein degradation (26S proteasome) (RPT1A, and RPT2A) **(Figure 1F, Supplementary Figure 5, and Supplementary Table 1)** ^29,30^, reinforcing the idea that SGs act as dynamic condensates integrating multiple biological processes.

During the recovery phase, we identified 95 proteins that collectively may orchestrate a return to normal cellular growth and function. These proteins include those involved in repairing damaged structures (CLPP6, RH40, RH12, RH14), strengthening cell walls (KCR1, UXS3, GPAT4), promoting growth and development (ML1, AHA2, CYT1), ROS-detoxification (PER69, PER71), and reinitiating translation (RACK1C, RPS2B, RPL8C, RPS27A, RPL9B) **(Supplementary Table 1)** ^31–41^. Furthermore, shared components between the heat and recovery phases highlight a link between ethylene and the jasmonic acid pathway (EIN2 and PLDBETA1) **(Supplementary Table 1)** ^42,43^, underscoring a coordinated reorganization of cellular processes during stress adaptation.

Previous studies have shown that salicylic acid (SA)-induced SGs contain several ribosomal proteins (r-proteins), linking the role of SGs to translational regulation under biotic related-stress conditions ^14^. Similarly, our study identified a robust cluster of r-proteins and translation-associated factors under all tested conditions **(Figure 1F and Supplementary Figure 5)**. However, the composition of r-proteins differed between our tested conditions, suggesting stress-specific ribosomal remodeling. To explore whether these differences were random or functionally coordinated, we mapped the distribution of the enriched r-proteins onto the ribosome structure using the ComplexOme-Structural Network Interpreter method (COSNet_i_) ^44^. Strikingly, these proteins were not randomly scattered but all clustered within the structure of the 40S small subunit (SSU) **(Figure 2A)**. Under recovery, new sets of r-proteins were enriched including also proteins from 60S big subunit, suggesting a targeted remodeling or recycling program in response to stress ^45,46^.

**Figure 2.**
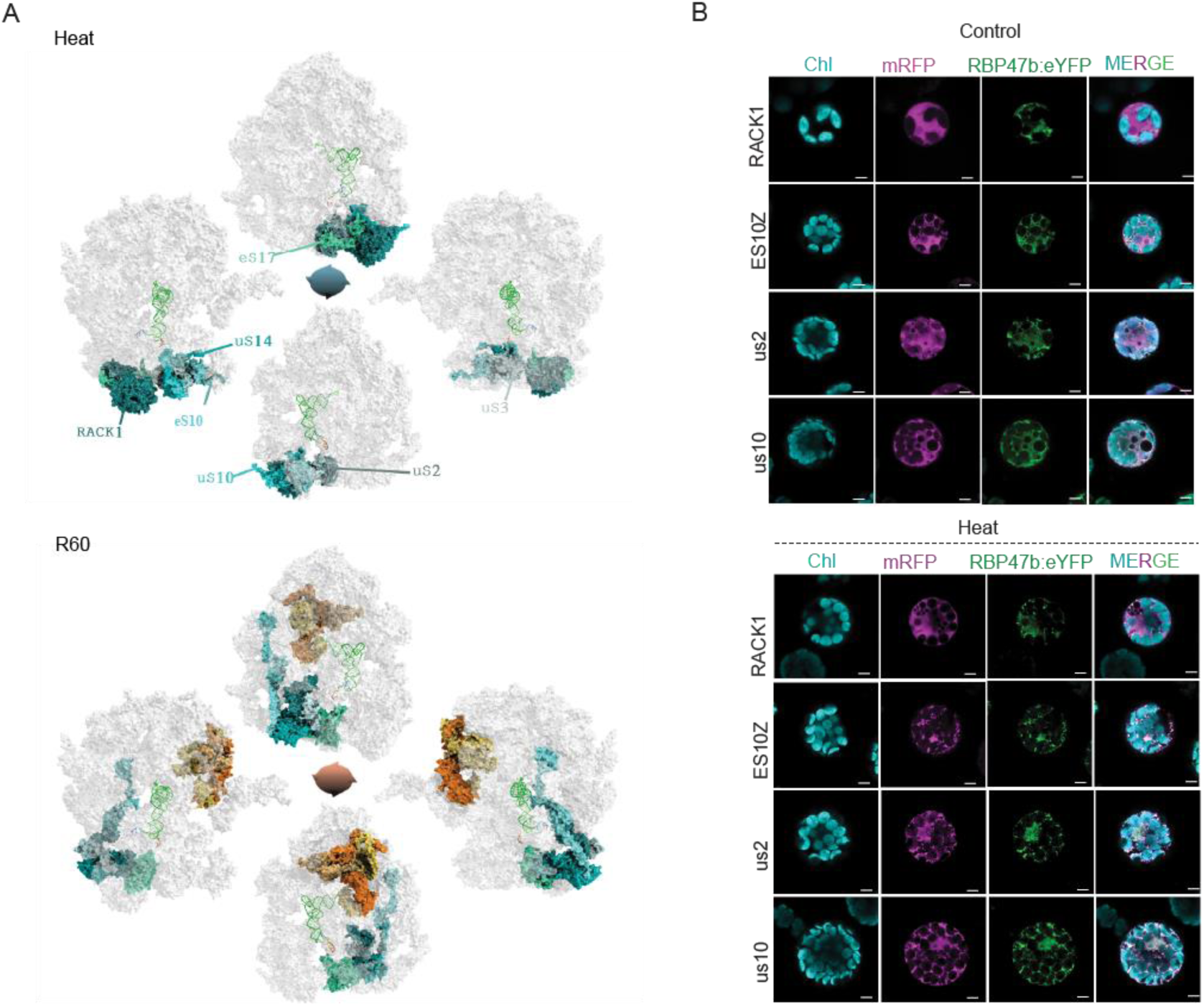
Proximity mapping of RBP47b SG proteins reveals a novel link to translation. **A**. Ribosomal proteins enriched in the interactome of RBP47b are structurally confined within the 40S small subunit. Spatial mapping of ribosomal proteins under heat stress, including their paralogs, within a rotated ribosome structure (90° increments). Proteins significantly enriched in the RBP47b interactome (based on adjusted P-values) under heat (top panel) or recovery 60 min conditions (R60, bottom panel) are significantly localized to the highlighted, structurally coherent regions within a spatial-adjacency ribosome graph sampled using a random walk methodology. **B**. Co-localization of selected ribosomal protein candidates fused to mRFP (magenta) in transiently transformed Arabidopsis protoplasts expressing *UBQ10p:RBP47b::EYFP* (green) under control (23°C) or heat conditions (42°C for 30 min). From left to right: Chlorophyll (Chl), mRFP, eYFP, and merged channels. Scale bars = 5 μm.

To further validate whether the key r-proteins such as RACK1A, eS10 (RPS10A), uS2 (RPSaA), and uS10 (RPS20A) that were identified as potential proximity interactors localized to RBP47b-SGs or remain in close cytosolic proximity, we generated Arabidopsis protoplasts from the *UBQ10p:RBP47b::EYFP* line and transiently expressed each r-protein fused to mRFP. Our analysis revealed that under heat stress conditions, the eS10, uS2, and uS10 co-localized with RBP47b within SGs, whereas RACK1A remained cytosolic **(Figure 2B)**, reinforcing a strong link between SG dynamics, ribosome remodeling, and translational control during stress adaptation.

### The RBP47 protein family associates with SGs under heat stress, regulating the timely disassembly of cytosolic mRNA foci

In Arabidopsis, RBP47b belongs to a four-member gene family consisting of RBP47 a, b, c, and c’, sharing 76%-92% of protein sequence similarity (**Figure 3A**) and a characteristic domain distribution with the presence of three RNA-recognition motifs (RRM) and Prion-like domains (PrLD). Thus, the RBP47 proteins may play similar biological roles. Indeed, existing datasets have previously identified RBP47a and RBP47c as SGs components ^11,13,14^. Consistent with this, our heat-stress proxitome analysis revealed the presence of RBP47a, RBP47c, and RBP47c’. Furthermore, transient expression assays in Arabidopsis protoplasts confirmed that RBP47c and RBP47c’ proteins co-localize with RBP47b in SGs under heat stress **(Figure 3B);** however, neither RBP47c or c’ show a nuclear localization as observed in the case of RBP47b. At the transcriptional level, all four family members exhibited a similar response to heat stress in Col-0 plants **(Supplementary Figure 6)**, suggesting that they may have overlapping functions in the plant heat stress response.

**Figure 3.**
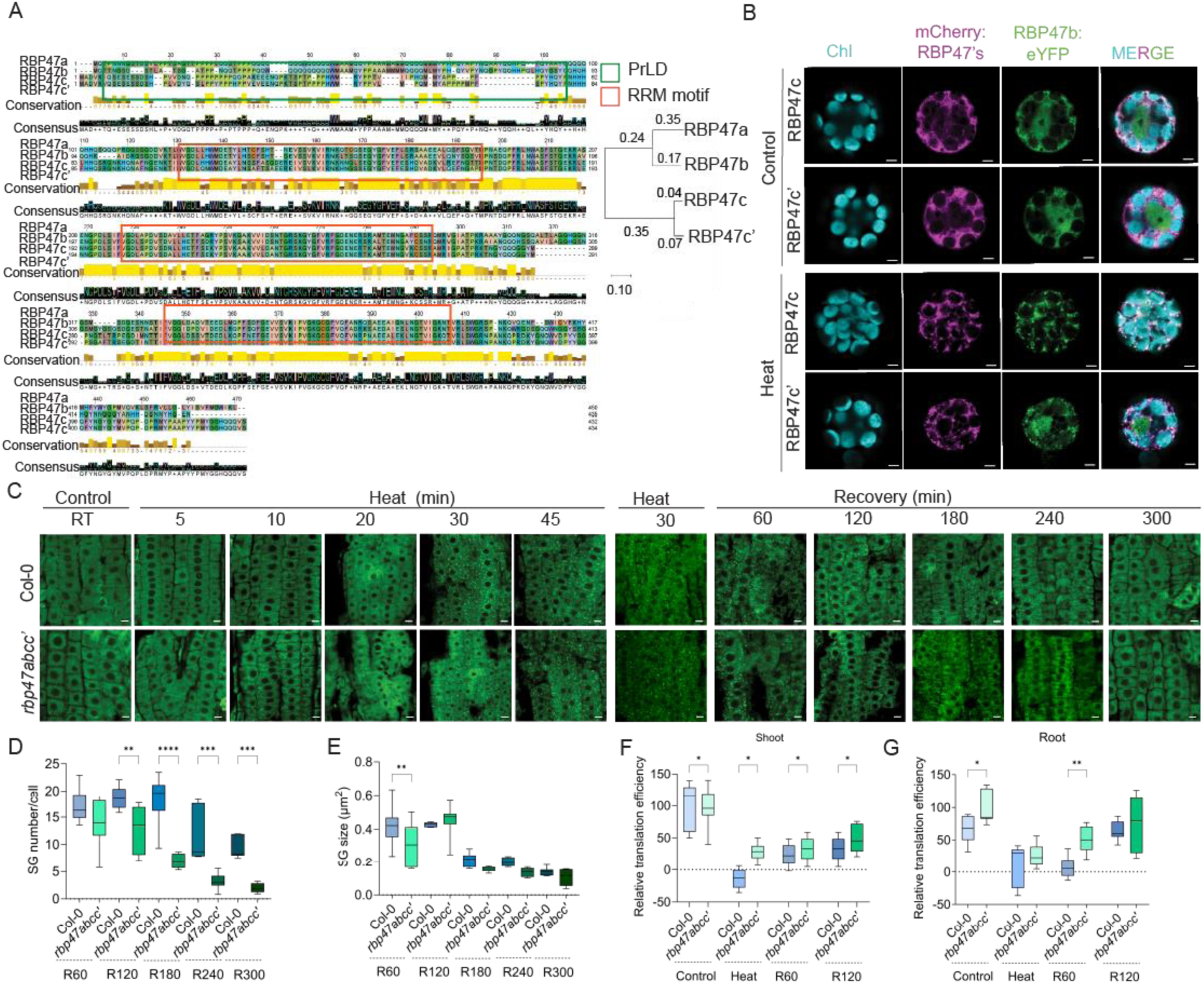
Proximity mapping of RBP47b SG proteins reveals a novel link to translation. **A**. Visualization of a multiple sequence alignment for the RBP47 family members. The alignment uses the BLOSUM62 substitution matrix to assign color codes ^48^. Conservation of the physicochemical properties of a given amino acid in each position is measured as a numerical index on a scale of 1-10. The consensus displayed below the alignment is the percentage of the modal residue per column. Green and orange boxes highlight conserved regions, such as PrLD and RRM motifs. The tree is drawn to scale, with branch lengths measured in the number of substitutions per site (next to the branches). **B.** subcellular localization of RBP47c and c’ fused to mRFP under control and heat conditions in transiently transfected protoplasts expressing RBP47b::YFP. *n* = 8, scale bar = 5 μm. **C**. poly(A)+ RNA FISH dynamics in Col-0 and *rbp47abcc’* exposed to control, heat, and recovery conditions. **D-E**. SGs number per cell and SGs size quantification (µm^2^) from panel **C**. Data is presented as mean ±SD n=6-, \**P* ≤0.05 and \*\*\*\**P* < 0.01; one-way ANOVA followed by Šídák’s multiple comparisons test. **F-G**. BONCATE, the relative translation efficiency in Col-0 and *rbp47abcc’* shoots and roots was evaluated at 42℃ (Heat) for 1h and left for recovery at 23℃ for 60 (R60) and 120 min (R120). Data represent the mean ± SEM, *n* = 3 biological replicates. \**P* ≤0.05 and \*\**P* < 0.01; one-way ANOVA followed by Dunnett’s multiple comparisons test.

To further explore the functional redundancy among RBP47 family members, we employed the previously reported *rbp47abcc’* mutant ^14^ **(Supplementary Figure 7A)** and evaluated its response to elevated temperatures. We first confirmed the reduced gene expression levels for all four RBP47 family members in the *rbp47abcc’* background (**Supplementary Figure 7B**). At the phenotypic level, 11-day-old Col-0 and *rbp47abcc’* seedlings growing under control conditions showed no differences in leaves (number and morphology) or root length **(Supplementary Figure 1B-C)**.

To determine whether the *rbp47abcc’* mutant retains the ability to form SGs under heat stress, we performed fluorescence in situ hybridization (FISH) using a Cy5-labeled oligo(dT) probe to detect polyadenylated RNA. FISH was carried out on Col-0 and *rbp47abcc’* seedlings under control, heat, and recovery conditions (**Figure 3C**). While poly(A)+ RNA showed a diffuse pattern under control conditions in both genotypes, the signal localized mainly into cytosolic foci under heat stress, confirming that both genotypes could form SGs in response to higher temperatures. When quantified, the *rbp47abcc’* mutant showed a slight delay in SG formation and, more interestingly **(Figure 3D-E)**, faster disassembly starting at 120 min compared to 240 min for Col-0 plants **(Figure 3C)**.

SGs are considered hubs for translationally stalled mRNA; their formation is associated with global translation inhibition ^47^. We hypothesized that the accelerated SG disassembly observed in *rbp47abcc’* mutants could lead to a more rapid restoration of the translational machinery. To understand whether global protein synthesis is different between Col-0 and *rbp47abcc’*, the biorthogonal non-canonical amino acid tagging combined with enzyme-linked immunosorbent assay (BONCATE) approach was employed ^14^. Shortly, in the BONCATE approach, a structural analog of methionine, azidohomoalanine (AHA), is incorporated into the newly synthesized proteins without inhibiting peptide chain elongation. The azido group on AHA allows subsequent covalent labeling of newly synthesized proteins with biotin–alkyne through the Click reaction, which can then be quantified in a high-throughput manner via ELISA.

To obtain a comprehensive view of how RBP47 proteins impact global protein synthesis, the BONCATE assay was performed on roots and shoots of Col-0 and *rbp47abcc’* plants exposed to control, heat, and two recovery time points (60 min and 120 min; **Figure 3F-G**). Our results confirmed that protein synthesis was inhibited in both genotypes upon exposure to heat stress conditions. However, striking differences emerged in recovery, where protein synthesis was restored faster in the *rbp47abcc’* mutants relative to Col-0 **(Figure 3F-G)**. The faster disassembly of mRNA foci and accelerated translational recovery in *rbp47abcc’* suggests that the RBP47 family plays a critical role in regulating the spatial and temporal organization of cytosolic mRNA foci, which might ultimately influence stress responses and tolerance.

### The *rbp47abcc′* plants reveal improved tolerance to acute heat stress conditions

Because our results implicated RBP47 proteins in heat response at the molecular level, we evaluated the *rbp47abcc’* plant phenotypic response to heat shock at the seedling stage. Briefly, 11-day-old Col-0, *rbp47abcc’,* and *hsp101* (sensitive to heat shock stress) ^49^ plants were subjected to 42°C for 5h, followed by 7 days of recovery (7 days after stress - 7-DAS) at 23°C **(Supplementary Figure 8A)**. As anticipated, all genotypes sustained severe damage following heat shock; however, the *rbp47abcc′* plants exhibited markedly superior recovery, evidenced by higher survival rates and greater chlorophyll retention (**Figure 4A-C**).

**Figure 4.**
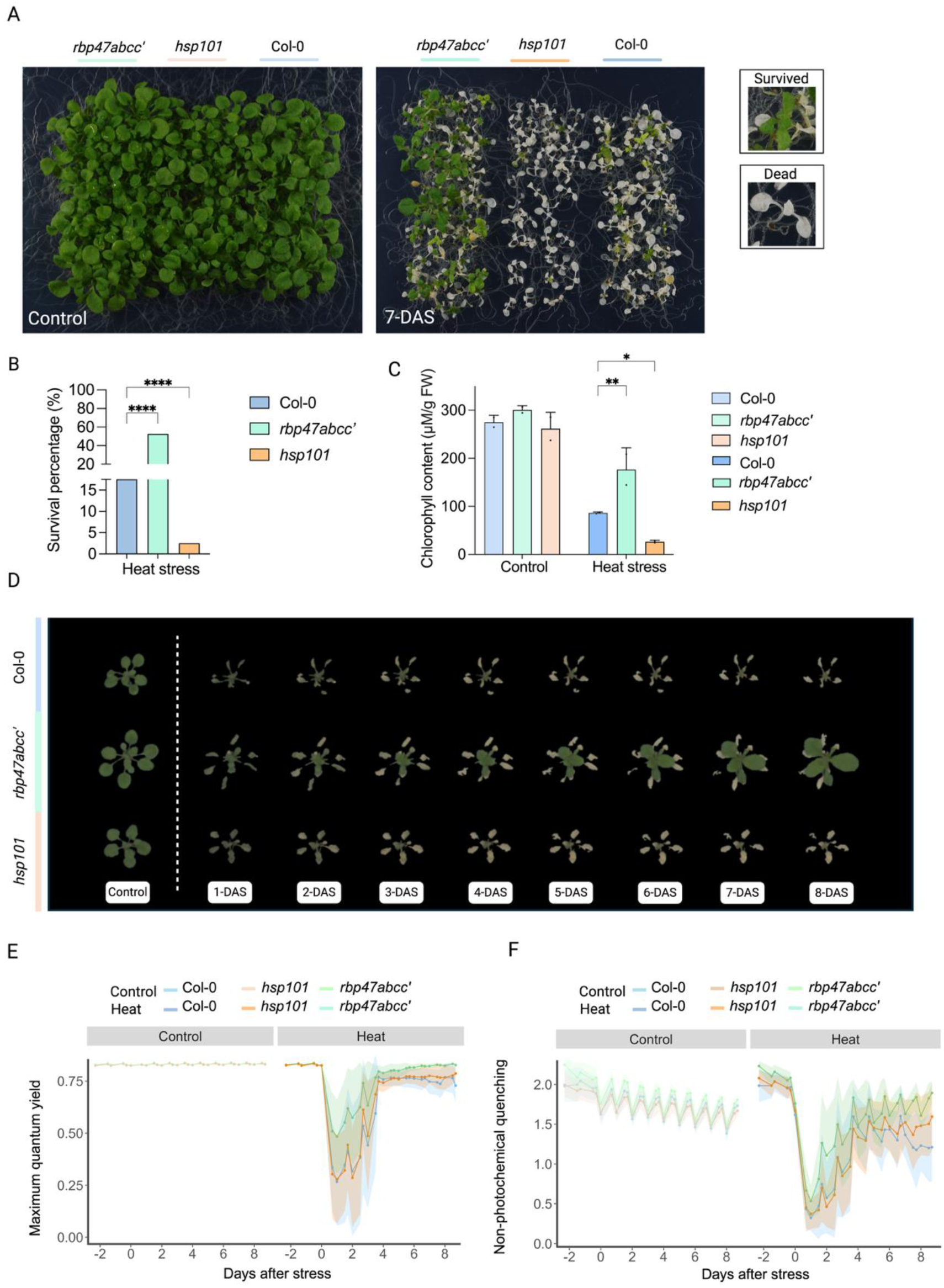
The *rbp47abcc’* mutant exhibits enhanced thermotolerance compared to Col-0 plants. **A.** Representative images of 18-day-old *rbp47abcc’*, *hsp101*, and Col-0 plants under control conditions (left) and 7 days after recovery from a 5h heat stress at 42°C (right). **B.** Survival rate of Col-0, *rbp47abcc’*, and *hsp101* plants following heat stress treatment. Data presented as % of total number of plants (n=3 plates, each plate with 20 plants per genotype). Data represented as mean ± SD, *P < 0.05; ** P < 0.005; **** P < 0.0001 (two-way ANOVA). **C.** Total chlorophyll content (µM/g FW) under control and heat stress conditions represented in the panel. **A.** Data represented as mean ± SD, *P < 0.05; ** P < 0.005; **** P < 0.0001 (two-way ANOVA). **D.** High-throughput time-course imaging of 22-day-old (Control) of Col-0, *rbp47abcc’*, and *hsp101* plants (n=50) under control and recovery for 8 days after heat stress (9 hours at 45°C). The images were captured daily at 00:30, starting from one day before stress to eight days after stress (DAS 8). **E.** Maximum quantum yield of PSII (Fv/Fm). **F.** NPQ values over the heat to the recovery period.

Intrigued by the increased heat stress tolerance observed at the early developmental stage, we challenged *rbp47abcc’* mutants at the rosette stage (22-day-old) for 9h at 45°C **(Supplementary Figure 8B)**. To evaluate its performance, we used an automated, noninvasive, high-throughput phenotyping facility (PSI, Photon Systems Instruments, Czech Republic) system that allowed us to monitor daily changes in plant morphology and photosynthetic performance ^50^. To eliminate positional bias, plant positions were randomized **(Supplementary Figure 8C)**. Under optimal growth conditions, all genotypes demonstrated similar developmental patterns, with consistent increases in rosette size and lack of notable differences in photosynthetic efficiency, indicating that in the absence of heat stress, *rbp47abcc’* does not intrinsically provide a growth advantage or disadvantage. However, after plants were challenged with 9h of acute heat stress, all genotypes showed immediate growth inhibition that was slowly restored in the *rbp47abcc’* mutant between 3- and 4-DAS during the recovery phase (**Figure 4D and Supplementary Figure 8E**). Canopy temperature measurements revealed that the *rbp47abcc’* plants dissipated heat more efficiently than the other genotypes **(Supplementary Figure 8D)**. Notably, 7-DAS, the *rbp47abcc’* mutants exhibited larger rosette sizes in contrast to Col-0 and *hsp101* plants **(Supplementary Figure 8E)**, suggesting that *rbp47abcc’* plants showed greater resilience to heat stress.

To understand the physiological basis for this improved growth recovery, we examined several photosynthetic parameters, including photosystem II (PSII) performance. As a consequence of heat stress, all genotypes demonstrated a decline in the efficiency of PSII photochemistry (QY max) (**Figure 4E**). However, the *rbp47abcc’* mutant was found to be less affected. The PSII quantum yield at its highest level in a dark-adapted state (QY_max) of PSII photochemistry revealed a physiological explanation for the improved performance of the *rbp47abcc’* background. 2-DAS, the QY_max value in *rbp47abcc’* plants stabilized at 0.62 ± 0.03, whereas in both Col-0 and *hsp101*, it decreased to < 0.50 (**Figure 4E**). These data suggest that the PSII machinery remained less affected and, therefore, more resilient in *rbp47abcc’*, allowing higher retention of photochemical efficiency compared to Col-0 and *hsp101*. This notion of the *rbp47abcc’* plants being more thermotolerant at the molecular level was also supported by the non-photochemical quenching (NPQ) values, which showed rapid restoration of NPQ capacity during the first days of recovery after stress (**Figure 4F**).

Furthermore, the electron transport rate (ETR), an indicator of the efficiency of photosynthetic electron flow, was significantly higher in *rbp47abcc’* plants throughout the recovery period, further supporting superior photosynthetic activity maintenance under heat stress **(Supplementary Figure 8F)**. Lastly, the chlorophyll retention and overall canopy vitality were analyzed by color segmentation, which showed a higher proportion of green tissue (RGB: 98,103,73) in the *rbp47abcc’* under heat conditions, whereas Col-0 and *hsp101* retained more brown tissue (RGB: 80,70,56), indicative of a damage **(Supplementary Figure 8G)**. Altogether, the data suggests that the RBP47 family may play a negative role in plant thermotolerance mechanisms.

### Transcriptomic data of the *rbp47abcc’* mutant revealed altered capacities for photosynthesis and hormone signaling

To better understand the molecular mechanisms underlying the enhanced thermotolerance of the *rbp47abcc’* mutant, we utilized 11-day-old plants and a multi-omics approach **(Supplementary Figure 9)** to examine their transcriptome, proteome, and metabolome compared to those of Col-0 plants under control, heat stress, and the recovery phase (60 min) **(Supplementary Figure 10)**. In Pearson correlation and principal component analyses of RNA expression profiles, Col-0 and *rbp47abcc’* samples under control conditions were strongly correlated and clustered separately from the heat and recovery samples of both genotypes **(Supplementary Figure 10A)**, reflecting their similar phenotypes under normal growth conditions. Under heat and recovery conditions, Col-0 and *rbp47abcc’* samples clustered separately from each other, suggesting distinct transcriptional responses to heat stress and the recovery phase.

Exposure to heat stress-induced widespread changes in transcript abundance in both genotypes, with 906 upregulated and 2,070 downregulated genes in Col-0 and 883 upregulated and 1,669 downregulated in *rbp47abcc’* **(Supplementary Table 2)** (differentially abundant genes, DAGs, defined as those with |log2FC| >1 and adjusted p-value <0.05). Substantial overlap was observed between Col-0 and *rbp47abcc’* DAGs **(Figure 5A)**. In both genotypes under heat stress, genes associated with heat response were enriched in the pool of upregulated transcripts **(Supplementary Figure 10B)**, demonstrating that loss of all four RBP47 genes does not impact the plant’s ability to perceive heat stress. Genes that were upregulated only in *rbp47abcc’* heat response were enriched for photosynthesis-related GO terms, and a suite of photosynthesis-associated genes (n = 26), regardless of DAG status, were upregulated in the mutant and downregulated in Col-0 in response to heat **(Figure 5B, quadrant 1)**. Enrichment of photosynthesis-associated genes was also observed when directly comparing the two genotypes under heat stress **(Supplementary Figure 10C)**. This increased abundance of photosynthesis genes under heat stress may underline the improved photosynthetic capacity of *rbp47abcc’* plants observed in high-throughput experiment on mature plants (**Figure 4**).

**Figure 5.**
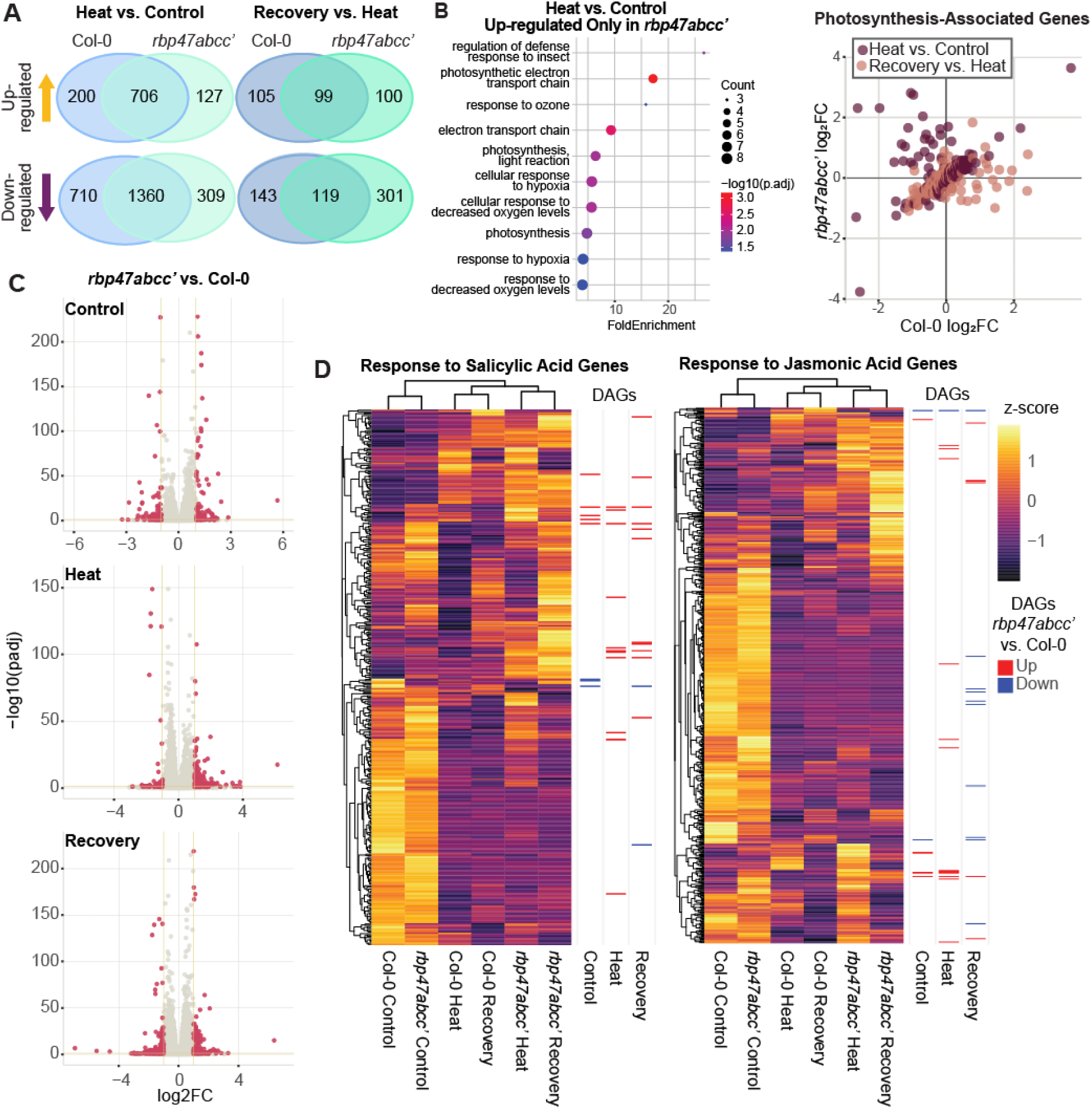
Examining the *rbp47abcc’*-specific response to heat stress and recovery conditions. **A.** Venn diagram of genes either up or downregulated in the two backgrounds upon the control to heat (left) or heat to recovery transitions (right). **B**. GO-enrichment of genes that are specifically up-regulated in the mutant background in heat stress conditions (left), with a comparison of transcript abundance changes (log2 fold change) between the two backgrounds for all genes with photosynthesis association (right). **C.** Comparison of differentially abundant transcripts in *rbp47abcc’* vs Col-0 under control, heat and recovery conditions. **D.** Heatmap depicting the expression variation of genes associated with either the SA (left) or JA (right) response under different conditions. Note, columns are clustered by similarity of response (hierarchical). To the right of each heatmap are markers denoting whether those genes were found to be significantly differentially abundant between *rbp47abcc’* vs Col-0 backgrounds in the three conditions.

Differential expression analysis between *rbp47abcc’* and Col-0 seedlings under each of the three conditions exposed small subsets of DAGs (**Figure 5C**; 79 up- and 83 down-regulated in control, 135 up- and 50 down-regulated in heat, 99 up- and 166 down-regulated in recovery) **(Supplementary Table 2)**. The majority of *rbp47abcc’* vs. Col-0 DAGs are unique to a single condition **(Supplementary Figure 10D)**, suggesting that loss of the RBP47s has context-dependent impacts on the transcriptome. Interestingly, GO enrichment analysis of DAGs revealed a significant number of genes associated with salicylic acid response to be upregulated in *rbp47abcc’* seedlings during both heat stress and recovery **(Supplementary Figure 10C)**. This pathway has been implicated in responses to myriad stresses ^51^. When examining all genes associated with salicylic acid response, *rbp47abcc’* heat and recovery samples clustered separately from Col-0 **(Figure 5D)**, indicating that this signaling pathway is regulated in a distinct way in the *rbp47abcc’* background under these conditions. Similarly, genes associated with response to JA showed a distinct expression profile in the *rbp47abcc’* mutant under heat stress and in recovery compared to Col-0 **(Figure 5D)**, with several DAGs under each of the conditions. Overall, transcriptome analysis of *rbp47abcc’* mutant suggested that altered capacities for photosynthesis and hormone signaling may support their improved heat stress resilience and that RBP47 family members may indirectly regulate these pathways under stress.

### RBP47 family is needed to direct reductions of global protein abundances towards specific parts of the proteome during heat stress

To understand the effect of the *rbp47abcc’* mutation on the status of active protein pools, we performed shotgun proteomics of whole plant tissue and compared total and individual protein levels with Col-0 **(Figure 6A and Supplementary Table 3)**. Overall, compared to its control, i.e., considering values of total protein accumulated relative to their own genotype control, the *rbp47abcc’* mutant had significantly more total protein during heat stress (fold change as compared to its control: 1.77E+08) and after 60 min of recovery from the heat stress stimulus (fold change as compared to its control: 1.14E+08) **(Figure 6B)**. In contrast, total protein decreased dramatically in Col-0 plants exposed to heat stress (fold change as compared to its control: -1.48E+07) and the recovery period (fold change as compared to its control: -1.92E+07) **(Figure 6B)**, suggesting that heat stress triggers a decrease in translation that is directly or indirectly dependent on RBP47. Thus, the *rbp47abcc’* mutant does not decrease protein contents as Col-0. Interestingly, the significant reductions in protein pools were constrained to specific biological functions and gene ontology terms. We found three major clusters using k-means hierarchical clustering based on an auto-scaled matrix and Pearson correlation as the distance metric, plus a bootstrap value of 1000 to ensure that the data fully supported clusters. From these three clusters (3, 4, and 5 in **Figure 6C**), the proteins that had significantly decreased in Col-0 plants but NOT in the mutant are associated with translation, ribosome, and photosynthesis (**Figure 6C**), suggesting that the *rbp47abcc’* mutant is translationally and photochemically more competent than the Col-0 under stress. The *rbp47abcc’* mutant also showed a greater accumulation of proteins related to SGs and photosynthesis (Cluster 3 & 4) **(Figure 6C)**. In this context, 15 out of 52 proteins from cluster 4 belong to the group of proteins essential for photosynthesis machinery, such as psa A-B, psb A-D, and light-harvesting complex (LHC) proteins. Indeed, when looking at the overall proteome profile, the three *rbp47abcc’* samples clustered with Col-0 recovery **(Figure 6C)**, suggesting that the mutant is not subjected to a full heat acclimation dynamic at the proteome level like the Col-0 but rather the *rbp47abcc’* mutant never acclimates, as the wild-type does. Interestingly, although heat response-associated proteins were not enriched in any cluster, *rbp47abcc’* was shown to specifically accumulate heat shock proteins such HSP70-4, HSP90-1, and HSP60-3 already under control conditions, further supporting the hypothesis that *rbp47abcc’* mutant is pre-acclimated at the proteome level.

**Figure 6.**
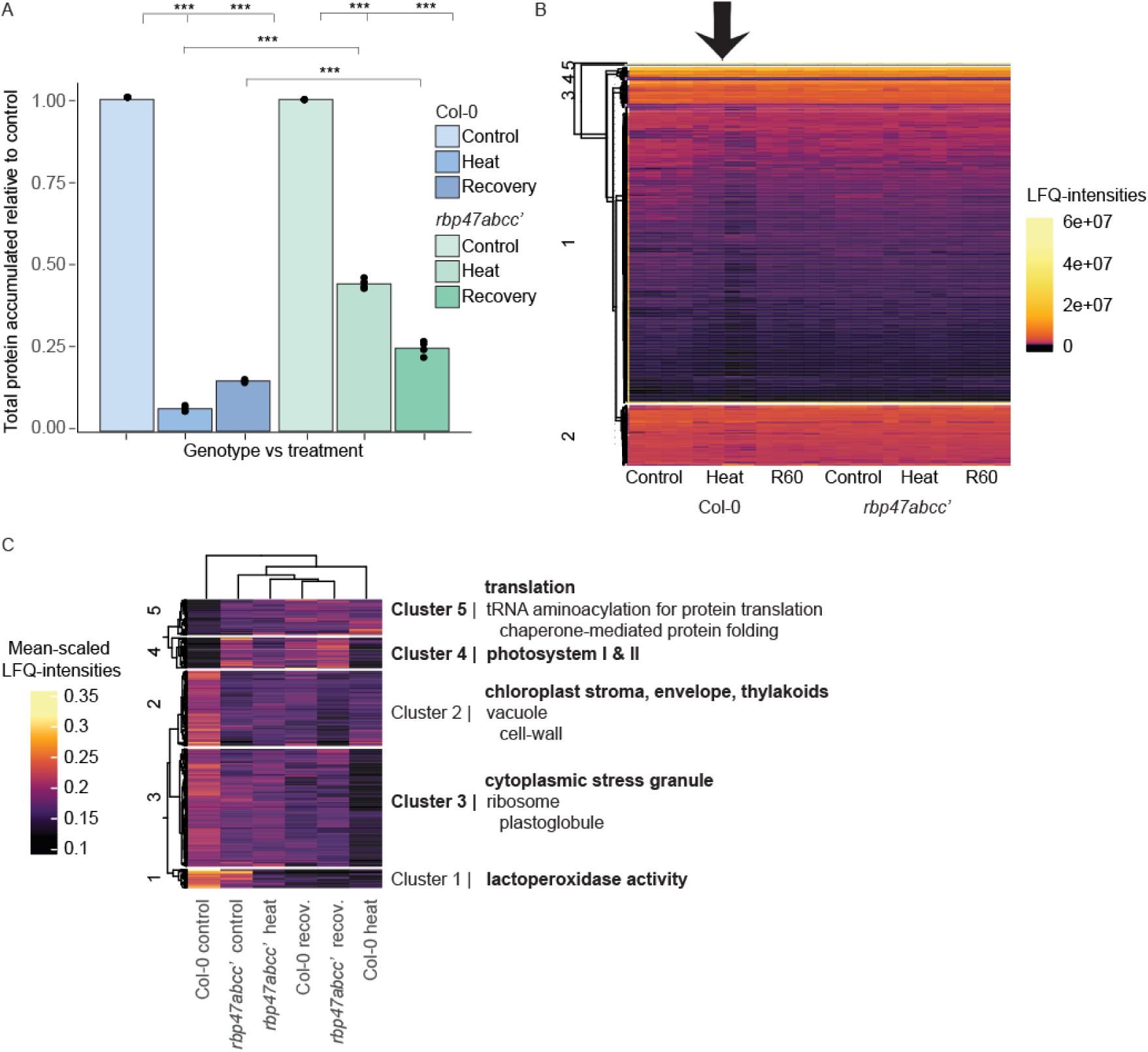
Untargeted statistical analyses of shotgun proteomics assays on Col-0 and *rbp47abcc’* plants during heat stress and a recovery period of 60 minutes. A. Barplot featuring total protein contents normalized to the non-triggered control sample across treatments. Total protein contents and statistical comparisons were made using pairwise Student’s t-tests with significance values (P) on the sum of label-free quantified (LFQ-normalized) protein abundances across four biological replicates (n = 4, *** means P < 0.01). **B**. Heatmap representation of all individual proteins detected using LC-MS/MS shotgun proteomics. Rows are LFQ abundances of individual proteins, and as such, k-partitions (from K-means = 5) represent groups of similarly abundant proteins in plant tissue. **C.** Heat map of scaled LFQ-abundances per individual protein. Columns are clustered using hierarchical clustering with Pearson correlation as the distance metric and complete as the clustering method. At the right side of each cluster, gene ontology terms (https://www.geneontology.org/) are significantly enriched in each protein subset.

### Metabolomics and lipidomics approaches suggested that the *rbp47abcc’* mutant accumulates essential molecules for plant stress detoxification

To identify changes in metabolic pathways that might explain *rbp47abcc’* heat tolerance, we analyzed the metabolic profile in 11-day-old Col-0 and *rbp47abcc’* under control, heat stress, and recovery phase (R60) using an untargeted and targeted metabolomics approach. All data sets were analyzed in MetaboAnalyst 6.0 software ^52^ to identify differentially accumulated compounds in the *rbp47abcc’* mutant under all tested conditions. A T-test (p-value ≤ 0.05, FDR) and fold-change analysis (≥ 2) were performed for pairwise comparison of both genotypes under the same conditions. Volcano plots from comparisons at control and heat conditions are shown in **Figure 7**, while a complete list of significant accumulated compounds can be found in **Supplementary Table 4.**

**Figure 7.**
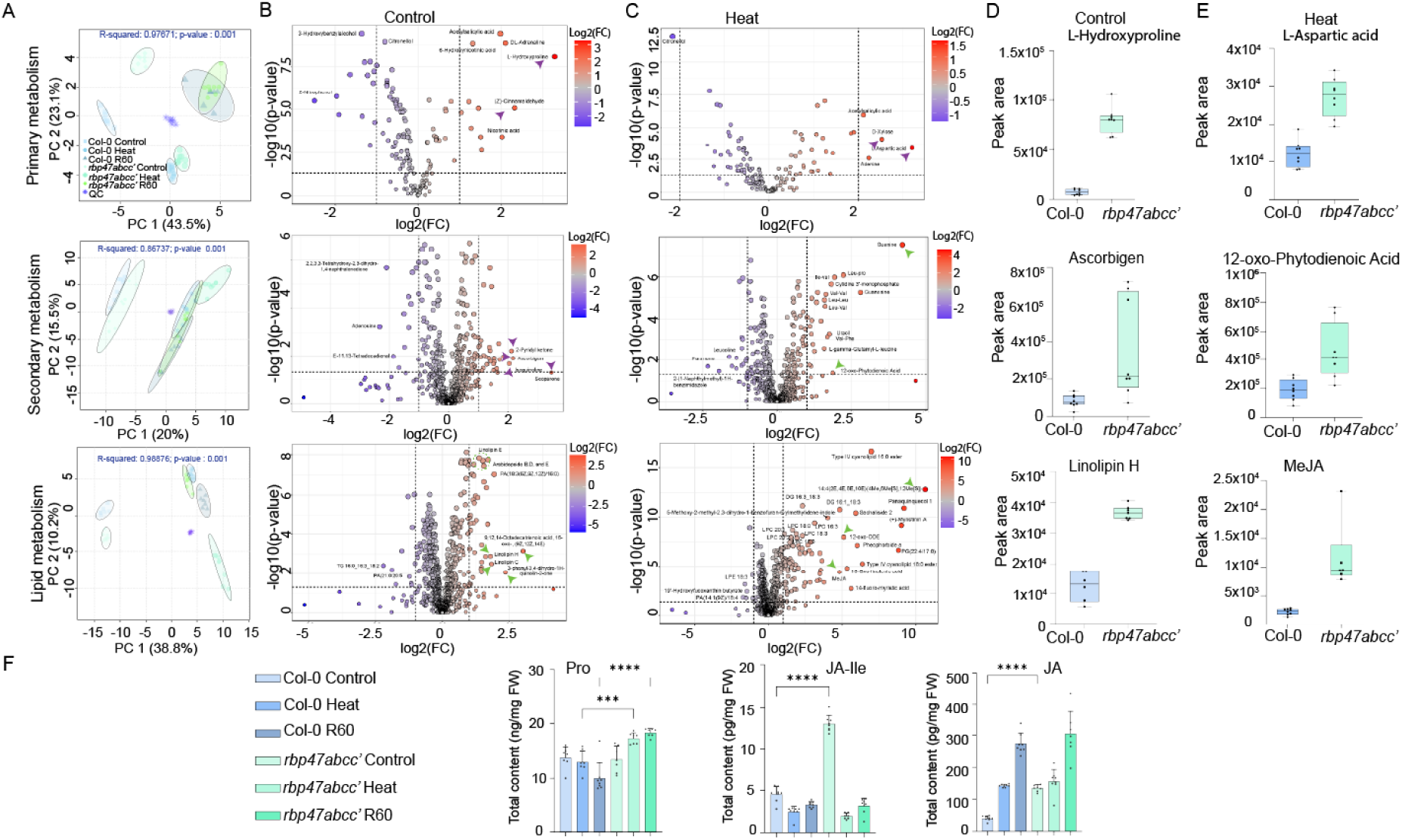
Metabolomics profiling reveals enhanced detoxification pathways in the *rbp47abcc’* mutant under control and heat stress. Differential metabolites and lipids identification in *rbp47abcc’* vs. Col-0 under control and heat conditions, datasets from primary metabolism are shown in the top panel, secondary metabolism middle panel, and lipid metabolism, bottom panel. **A**. Principal component analysis (PCA) score plots of 8 replicates for primary (GC-MS, top panel), secondary (middle panel), and lipid (LC-MS, bottom panel) untargeted data sets. The statistical significance of the group separation in PCA is evaluated using PERMANOVA, and distributions are computed using the Euclidean distance based on the PCs (R-square) and the overall significance (p-value). **B-C**. Volcano plot analysis showing differentially accumulated metabolites (DAMs) in *rbp47abcc’* vs. Col-0 under control **B** and heat stress **C.** Metabolites related to detoxification (purple arrows) and the Jasmonic acid pathway (green arrows and dashed lines) are highlighted. **D-E**. Boxplots showing the relative abundance (peak area) of representative metabolites extracted from **B-C** panels under control (**D)** and heat stress (**E)** conditions. **F**. Targeted metabolomics analysis of Proline (Pro), Jasmonic Acid (JA), and Jasmonate-Isoleucine (JA-Ile) in Col-0 and *rbp47abcc’* under control, heat stress, and recovery conditions. Data are represented as mean ± SD (n = 6–8). Statistical significance was assessed using one-way ANOVA with Šídák’s multiple comparisons test (*p < 0.05, **p < 0.01, ***p < 0.001, \*\*\**p < 0.0001*).

To this effect, we first explored differences in the primary metabolism by analyzing GC-MS profiles of both genotypes under all conditions tested. A total of 510 compounds were identified. 183 unknown compounds and 172 compounds with low group variance (≤20%) were removed. A total of 156 known compounds were included in the data analysis. The principal component analysis (PCA) of these compounds allowed the characterization of samples according to tested conditions and genotypes **(Figure 7A, top panel)**. The separation along PC1 (43.5%) vs PC2 (23.1%) suggests that condition-specific effects drive the primary source of variation. The clustering of the samples between genotypes and across conditions indicated a differential primary metabolic response of *rbp47abcc’* and Col-0 under control conditions, heat stress, and recovery phase (R60) (**Figure 7A, top panel**).

Under control conditions, the *rbp47abcc’* mutant preferentially accumulated primary metabolites such as the L-Hydroxyproline, (Z)-Cinnamaldehyde, and Nicotinic acid **(Figure 7B and 7D)**. Interestingly, these compounds have been reported to play a role in plant oxidative stress responses ^53–56^. On the other hand, the down-regulation of phenolic compounds in *rbp47abcc’,* such as the 2-Nitrophenol ^57^, suggested that the plant is better prepared to face the stress without a constitutive activation of intensive detoxification mechanisms **(Figure 7B, top panel)**. Under heat conditions, minimal changes between genotypes were detected **(Figure 7C and E, top panel, and Supplementary Table 4).**

As a part of the untargeted approach, we also explored the secondary metabolite profile. Overall, 1,325 compounds were identified, of which 337 unknown compounds were removed, and 247 compounds were filtered out based on group variance (≤ 20%). A total of 741 compounds were thus included in the data analysis. As observed, the PCA clearly separated the genotypes between control and heat conditions. The separation along PC1 (20%) vs PC2 (15.5%) suggests that the primary source of variation is driven by condition-specific effects **(Figure 7A, middle panel)**. The clustering of the samples under each condition suggests a differential activation of metabolic pathways or damage repair mechanisms among genotypes. Under control conditions, the *rbp47abcc’* mutant showed a significant differential accumulation of compounds related to cellular redox homeostasis, such as Scoparone (a derivative of the coumarins), Isoquinoline, Ascorbigen, L-gamma-Glutamyl-L-leucine **(Figure 7B, middle panel)** ^58–62^.

Under heat stress, a suite of molecules related to nucleic acid synthesis, salvage pathways, and amino acid recycling (Guanine, Guanosine, Uracil, and Cytidine 3’-monophosphate, Leu-Pro, Ile-Val, Val-Phe, Val-Val, Leu-Leu, Leu-Val) was observed **(Figure 7C, middle panel)**^63^. Noticeably, the targeted analysis profile of amino acid in the *rbp47abcc’* mutant revealed a clear trend to accumulate higher levels of free amino acids after 60 min of recovery (R60) **(Supplementary Figure 11A)**. Notably, proline, a well-known reactive oxygen species (ROS) scavenger, showed higher accumulation in the *rbp47abcc’* mutant compared to Col-0 under heat and recovery (R60) conditions **(Figure 7F)**. Another interesting finding was the accumulation of the 12-oxo-phytodienoic acid (OPDA) in the *rbp47abcc’* plants. OPDA is a primary precursor in JA biosynthesis and has been shown to mediate adaptive responses to various forms of biotic and abiotic stresses ^64^ **(Figure 7B-E, middle panel)**.

The most striking differences observed in the *rbp47abcc’* mutant was observed in the lipid profile under both control and heat conditions. Of the 5,617 detected compounds, 4597 were unidentified, and 408 were removed based on group variance, leaving 610 compounds to be included in the analysis. Principal component analysis (PCA) of lipidomics data revealed a clear clustering pattern driven by stress treatment and genotype. PC1 (38.8% of variance) primarily separated samples by treatment, while PC2 (10.2%) distinguished the two genotypes. Heat stress caused a pronounced shift in lipid profiles away from the control cluster. Overall, both genotypes showed a similar response to heat, but they remain separated from each other, suggesting underlying differences in their profiles and stress-induced lipid changes **(Figure 7A, bottom panel)**.

Under control conditions, the *rbp47abcc’* mutant accumulated higher levels of stress-related lipids relative to Col-0. Notably, oxylipin-containing compounds such as arabidopside B and D and linolipins C and H were elevated, as was the 16-oxo-9, 12, 14-octadecatrienoic acid (16-oxo-OTA), an oxidized fatty acid derivative **(Figure 7B, bottom panel)**. These molecular species are typically formed under heat stress ^65^. In addition, the *rbp47abcc’* mutant showed higher basal levels of phosphatidic acid (PA 18:3/16:0), a well-known lipid stress-related second messenger in plants ^66^. Elevated oxylipin and PA levels in the absence of stress indicate that the mutant maintains an enhanced signaling readiness.

Upon heat stress, key membrane structural lipids showed changes indicative of adaptation to high temperatures, such as digalactosyldiacylglycerol (DGDG) species and monogalactosyldiacylglycerol breakdown products (MGMG 18:3) **(Figure 7C, bottom panel and Supplementary Table 1)**. An increased DGDG:MGDG ratio is known to preserve thylakoid membrane integrity under heat ^67,68^. Beyond structural lipids, the *rbp47abcc’* mutant uniquely accumulated a suite of stress-signaling molecules and protective metabolites. Notably, it showed elevated levels of methyl jasmonate (MeJA, methyl epoxy-octadecadienoate) **(Figure 7C-D)** and an oxo-fatty acid (9-oxo-10,12-octadecadienoic acid), both of which are potent jasmonate-family oxylipins involved in orchestrating stress responses ^68^. Since both secondary metabolite and lipid profiles indicate an accumulation of JA precursor and derivatives, we decided to explore hormonal alteration in a targeted approach by measuring jasmonic acid (JA), jasmonate-isoleucine (JA-Ile), salicylic acid (SA), indole-3-acetic acid (IAA), and abscisic acid (ABA) **(Figure 7F and Supplementary Figure 11B)**. As expected, total JA content and JA-Ile were higher in the *rbp47abcc’* mutant than in Col-0 under basal conditions **(Figure 7F)**. Interestingly, Col-0 accumulates more SA under control and recovery conditions. Other analyzed hormones were not significantly enriched in *rbp47abcc’* mutant **(Supplementary Figure 11B)**.

In parallel, under heat stress, the *rbp47abcc’* mutant accumulated higher chlorophyll catabolite sub-products such as pheophorbide A ^69,70^. Interestingly, we found higher chlorophyll content in the *rbp47abcc’* mutant compared to Col-0 after 60 min recovery from heat stress **(Supplementary Figure 11C)**, suggesting an accelerated chlorophyll-turnover cycle, where *rbp47abcc’* channels damaged chlorophyll into the catabolic pathway earlier under heat and rebuilds faster the photosynthetic antenna once the stress is removed. On the other hand, we also found an increased accumulation of antioxidant-related compounds such as vitamin E (α-Tocopherol) derivatives, which have been shown to prevent photo-oxidative damage by dismantling potentially harmful light-harvesting pigments, and membrane peroxidation under heat stress ^71^. Glucosinolates and carotenoids were also measured, but no statistical differences were found between genotypes and among treatments **(Supplementary Figure 11C and Supplementary Figure 12)**

Overall, our data pointed to a basal elevation of stress-associated compounds in the *rbp47abcc’* mutant that likely confers a primed state, enabling faster sensing and response to heat stress. This agrees with transcriptomic and proteomic data, where *rbp47abcc’* was pre-acclimated to stress. Under acute heat stress, the amplified production of antioxidant and stress-relief compounds, as well as the extensive lipid profile remodeling, helps to neutralize ROS and prevent heat-induced oxidative injury.

### The *rbp47abcc’* accumulates less ROS in comparison to Col-0 plants

Based on the clear indication that *rbp47abcc’* plants exhibit enhanced recovery from high-temperature stress, probably due to improved photosynthetic efficiency and detoxification capacity, we thought it crucial to examine the ROS levels in both genotypes. Additionally, as JA has been implicated in mitigating oxidative stress ^72^ and our metabolomics data revealed increased levels of JA and its derivatives in the *rbp47abcc’* mutants, we investigated the effect of exogenous application of MeJA on ROS accumulation. To this aim, we employed 3,3′-diaminobenzidine (DAB, Sigma) and 2′,7′-dichlorofluorescein diacetate (DCFH-DA, Sigma) staining techniques to detect H_2_O_2_ and ROS *in situ,* in 7-day-old Col-0 and *rbp47abcc’* plants under control and heat conditions.

Leaves of both genotypes did not show differences in the accumulation of H_2_O_2_ under control and heat conditions **(Figure 8A).** Interestingly, when analyzing roots, Col-0 plants accumulated more H_2_O_2_ and ROS under control conditions in comparison to *rbp47abcc’* plants. The accumulation was pronounced, particularly in the meristematic and elongation zones of the primary root **(Figure 8A-B)**. Relative quantification of ROS levels revealed that under control conditions, *rbp47abcc’* plants accumulate ∼30% less ROS than Col-0. Under heat stress, ROS accumulation in Col-0 increased by ∼25%; meanwhile, the *rbp47abcc’* plants showed an increase of less than ∼10%. **(Figure 8C).** The most spectacular ROS reduction effect was observed under the exogenous application of 5 µM of MeJA, where both genotypes exhibited a reduction of ∼70% **(Figure 8C)**. These results reinforce the hypothesis that elevated concentrations of JAs in the *rbp47abcc’* plants contribute to their improved ROS scavenging capacity under stress.

**Figure 8.**
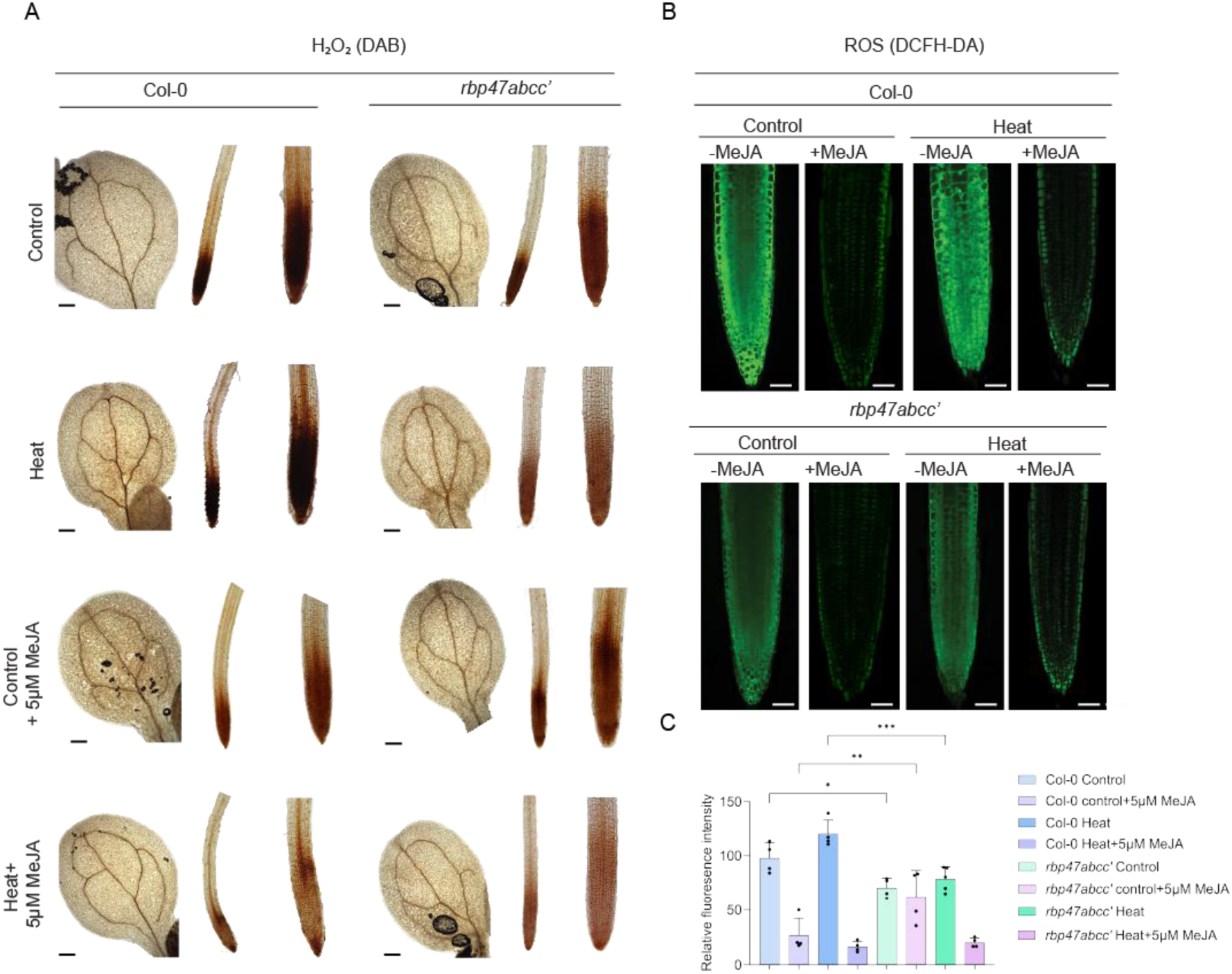
The *rbp47abcc’* mutant accumulates lower levels of reactive oxygen species (ROS) than Col-0. **A.** Hydrogen peroxide (H₂O₂) accumulation, visualized by 3,3’-diaminobenzidine (DAB) staining, in cotyledons and primary roots of Col-0 and *rbp47abcc’* seedlings under control conditions (23°C) and 1h of heat stress (42°C), as well as after treatment with 5 μM methyl jasmonate (MeJA) for 1h prior respective treatment. Scale bars = 200 μm. **B.** DCFH-DA staining for ROS in the primary root of Col-0 and *rbp47abcc’* under control, after heat treatment, and a combination of control or heat supplemented with 5μM MeJA for 1h prior to treatment. Scale bars = 50 μm. **C.** Relative staining intensity calculated from panel B using ImageJ software. Data is shown as the mean ± SD (*n* = 4). ^∗^*P* < 0.05; ^∗∗^*P* < 0.01; ^∗∗∗^*P* < 0.001 by Dunnett’s multiple test comparison. All the experiments have been repeated at least three times.

## Discussion

In the present study, we report for the first time a negative regulatory role of the RBP47 family in plant stress tolerance. Building on the selection of RBP47b as a *bona fide* SG marker ^13^, we adapted proximity labeling approach to capture RBP47b’s dynamic interactome under control, heat stress, and recovery conditions. This strategy was designed to gain deeper insights into the molecular remodeling of RBP47b-associated complexes during SG assembly, maintenance, and disassembly. To better understand the role of the RBP47 family in heat stress, we complemented this study with a multi-omic analysis in Col-0 and *rbp47abcc’* mutant backgrounds. Interestingly, we found that RBP47b -SGs are associated with proteins related to translation, photosynthesis, hormones, and detoxification pathways, suggesting a multifactorial thermotolerance response.

### RBP47b SGs Rewire the Translational Landscape

Here, we reported dynamic associations of RBP47b with specific r-proteins. Under control conditions, RBP47b was found to associate with a broad set of r-proteins. However, upon heat stress, these associations were dramatically restructured with RBP47b preferentially interacting with proteins belonging to the 40S small ribosomal subunit. Once the stress subsided, the interaction network remodeled yet remained predominantly linked to 40S components, indicating a stress-specific reorganization of ribosomal associations that likely underpins selective translation restart during recovery. In line with these findings, we observed a faster disassembly of poly(A)+ RNA foci in the *rbp47abcc’* mutant, which was linked to a faster translation recovery in the mutant compared to Col-0 plants in our BONCATE experiments.

Consistent with our findings, quantitative proteomics revealed that total protein abundance in *rbp47abcc’* plants showed a modest decline compared to Col-0 plants and sustained levels throughout recovery. A recent study showed that the RBP47 family influences ribosome sequestration into SGs upon SA treatment and is required for global translation inhibition ^14^. In our heat stress model, however, both Col-0 and *rbp47abcc’* exhibited translational repression, differing from the SA-induced response reported previously. Notably, the faster SG disassembly and enhanced thermotolerance observed in *rbp47abcc’* suggest that a faster restoration of global translation contributes to accelerated growth and recovery compared to Col-0 following heat stress.

One possible scenario is that RBP47 proteins play a role in 40S subunit remodeling, a role that allows for selective translation that promotes survival in stressful environments. In this line, a study in yeast demonstrated that the 40S subunit remodeling of preexisting ribosomes led to a rapid production of distinct ribosome populations to enable a translational response to high Na+, sorbitol, or pH stress ^46^. Similarly, a study in barley showed ribosome structural remodeling in the 60S and 40S subunits under cold conditions, allowing the cells to respond and adapt to cold conditions ^45^.

Our evidence indicates that the RBP47 family might be a part of a translational checkpoint, particularly during heat stress. By selectively sequestering 40S ribosomal subunits into the cores of SGs, RBP47 proteins might restrict subunit remodeling, thereby delaying the reactivation of protein synthesis. Likely, in the *rbp47abcc’* mutant, the 40S subunits are more available for rapid structural remodeling and ribosome reassembly, expediting translational recovery and conferring superior thermotolerance.

### Safeguarding Photosynthetic Capacity via Selective SG Docking

Photosynthesis is a highly temperature-sensitive process due to the detrimental effect on enzymes under elevated temperatures that lead to impairment of the PSII function, decrease in electron transport rates, inhibition of Rubisco activase, and decrease in the chlorophyll content ^73,74^. In all proxitome datasets, we detected the enrichment of photosynthesis-related proteins. The specific enrichment of core PSII/PSI subunits rather than the entire thylakoid proteome argues against random carry-over and supports a targeted interaction. These data prompted a physiological test of the *rbp47abcc′* mutant at early- and late-vegetative stages. In both cases, the mutant exhibited markedly greater thermotolerance than Col-0. The *rbp47abcc’* mutant retained more chlorophyll and survived better after acute heat stress. Meanwhile, mature rosettes showed higher PSII efficiency (Fv/Fm and NPQ) compared to Col-0 or the heat-shock-sensitive mutant *hsp101*.

Transcriptomic data revealed accumulation of photosynthesis-associated mRNAs in *rbp47abcc′* mutant under heat stress, and quantitative proteomics showed selective accumulation of core PSI/PSII and light-harvesting complex (LHC) under heat stress. Therefore we, propose that RBP47b -SGs bind a defined subset of photosynthetic mRNAs and nascent proteins during heat, transiently suppressing their translation or targeting them for turnover. In the absence of RBP47 proteins, these transcripts and protein fragments remain accessible, allowing faster translation and more rapid re-assembly of the photosynthetic apparatus once the stress subsides. Whether RBP47 proteins stabilize photosynthetic transcripts or, conversely, repress their expression remains unresolved; their absence clearly accelerated translational recovery and enhanced thermotolerance in *rbp47abcc′* plants.

Nevertheless, RBP47s are not chloroplast proteins. Recent studies suggest, that stress conditions might suppress the import of nascent proteins into chloroplast, therefore they might accumulate in the cytosol ^75^. Interestingly, a recent report showed that SGs can directly contribute to the cellular damage response by physically acting as molecular plugs to patch up leaky endomembranes in mammalian cells ^76^. This process was reported to involve ESCRT-dependent and independent pathways. Interestingly, in our proximity datasets, we also found components of ESCRT machinery. In this regard, it could be possible that RBP47b -SGs help to stabilize chloroplast outer membranes under heat conditions, though this hypothesis will require further testing. A plausible physical link could be the chloroplast-associated protein degradation (CHLORAD) pathway, which exports damaged outer-envelope proteins to the cytosolic 26S proteasome for its degradation ^77^. Although CHLORAD has not yet been shown to act on internal thylakoid proteins, our proximity list is enriched for HSP70-4, HSP90-1, HSP60-3, multiple 26S subunits, chloroplast ATPases, and TOC–TIC components, placing RBP47b SGs near chloroplast protein-quality-control hotspots during heat stress.

### JA-Primed Antioxidant and Detoxification Networks Limit ROS Build-Up

While the loss of RBP47 proteins expedited the restart of translation and reassembly/protection of the photosynthetic apparatus after heat stress, our proxitome and multi-omics analyses revealed that this remodeling extends into stress-hormone signaling and detoxification pathways. At the transcript level, a pronounced rewiring of SA and JA pathways was observed in the *rbp47abcc’* mutant under heat and recovery conditions. A similar trend was observed in our lipidomics profiles under heat, where a significant accumulation of stress-signaling lipids such as oxylipins, primary precursors of JA (OPDA), and JA-derivatives (MeJA) was observed in the *rbp47abcc’* mutant. Targeted hormone quantification confirmed that *rbp47abcc′* seedlings contain elevated basal levels of bioactive jasmonates (JA and JA-Ile) under control conditions, whereas Col-0 accumulates more SA during control and recovery phases. Notably, the antagonistic effect between the SA and JA defense pathways has been documented ^78^. Activation of the JA pathway suppresses SA biosynthesis and signaling, leading to a reduction in cellular ethylene sensitivity, thereby restricting programmed cell death ^78,79^. The elevated basal pool of bioactive JA observed in the *rbp47abcc′* mutant provides a practical explanation for the muted SA response during heat and recovery phases.

JA is well-known for its role in a wide range of biotic and abiotic stresses ^79,80^. Considerable evidence has supported the role of JA in alleviating intracellular ROS levels by activating genes coding for antioxidants and associated defense proteins ^79,81,82^. In line with these reports, the untargeted profiling of primary and secondary metabolites revealed that *rbp47abcc’* seedlings accumulate more molecules linked to oxidative stress protection and detoxification at baseline, including coumarin precursors ((Z)-Cinnamaldehyde) and derivatives (scoropone), alkaloids (isoquinoline), and glucosinolate breakdown products (ascorbigen) ^56,58–60,62,83^. In our targeted approach, we also detected high proline accumulation under heat and recovery conditions in the *rbp47abcc’* mutant. These molecules have been extensively reported to contribute to cellular redox homeostasis ^55^.

Together, these metabolite profiles indicate that the mutant is metabolically “on alert” for stress, maintaining an enhanced hormone signaling and antioxidant capacity from the outset. This was evidenced by the markedly lower accumulation of reactive oxygen species (ROS) in *rbp47abcc’* tissues at basal conditions. Mutant seedlings contained ∼30% less ROS than Col-0 even under control conditions, and unlike Col-0, the *rbp47abcc’* showed no significant spike in ROS after heat exposure. Notably, exogenous application of MeJA to Col-0 significantly reduced ROS levels by ∼70% under heat. This confirms that the heightened JA status of *rbp47abcc’* actively contributes to its superior ROS-scavenging capacity.

In this sense, SGs might offer an additional regulatory layer by transiently sequestering signaling nodes and enzymes, thereby imposing precise timing on pathway re-engagement, an idea demonstrated for TORC1 in mammals ^84^. Consistent with this model, all three RBP47b proximity datasets were enriched for detoxification, lipid metabolism, and JA biosynthesis enzymes. We propose that SG-mediated co-localization of these modules creates an integrative hub in which redox status, membrane-derived oxylipins, and jasmonate production are synchronized during heat stress. In the *rbp47abcc′* background, altered SG composition and faster granule disassembly likely accelerate the release—and hence the activity—of detoxification and JA-biosynthetic enzymes, shifting the hormonal balance toward JA and further suppressing SA output. This coordinated rewiring of hormone signaling and antioxidant capacity provides a mechanistic underpinning for the mutant’s superior control of ROS and enhanced thermotolerance.

## Conclusion

In summary, our findings establish the RBP47 family as critical regulators of SG dynamics, translational reprogramming, and metabolic adaptation during heat stress. Loss of RBP47 function primes plants for enhanced thermotolerance by accelerating translational recovery, stabilizing photosynthetic performance, and reinforcing antioxidant and hormone-mediated defense pathways. This pre-activated stress-resilient state underscores a broader role for RBP47 proteins in fine-tuning cellular homeostasis under adverse conditions. From an evolutionary aspect, it seems beneficial for plants to have RBP47 proteins present to avoid long-term activated “on alert” mechanisms. Thus, negative regulation by the RBP47 family might represent an evolutionary trade-off that allows plants to balance heat protection with metabolic economy in unpredictable environments. Future work dissecting the molecular interfaces between RBP47-associated condensates, translational machinery, and organelle quality control systems will be instrumental in elucidating new strategies to engineer stress-resilient crops.

## Material and methods

### Plant material and growth conditions

Col-0 was used as wild-type (WT) plants in this study. Homozygous CRISPR (Cas)9-free *rbp47abcc′* were obtained from Dr. Wei Wang (State Key Laboratory for Protein and Plant Gene Research, School of Life Sciences, Peking University, Beijing, China). For the present study, we generated *UBQ10p:RBP47b::eYFP/Col-0* and *UBQ10p:RBP47b::eYFP/rbp47abcc’* lines to study SGs dynamic and mRNA foci dynamics and *UBQ10p:RBP47b::mTb::NES::mVenus/Col-0* and *pUBQ10:mTb::mVenus/Col-0* transgenic lines for proximity labeling assay. The expression levels of all lines were confirmed (**Supplementary Figure 1**). Arabidopsis seeds were surface sterilized with bleach 20% for 3 min, followed by one wash with ethanol 70 %, triton X-100 0.05% for 3 min. The solution was removed, and the seeds were washed 5x with sterile MiliQ water and left at 4 °C in the dark for 48 h. Arabidopsis seeds were sowed on either 1/2 Murashige and Skoog (MS) (Sigma) plates with 1% (w/v) sucrose and 0.7% (w/v) agar or liquid half-strength MS media (for proximity labeling and ROS assays). For all experiments, the plants were grown at 23 °C in long-day conditions (16h light/8h dark with 120 μM m^−2^ s^−1^) for 11 days, unless otherwise stated.

### Construct generation

RBP47b (AT3G19130) DNA sequence without stop codon, flanked by AttL1 and AttL2 recombinant sites, was synthesized (GeneScript Biotech, Singapore). The UBQ10 promoter sequence was PCR amplified using as a template the Addgene vector #127369, using Phusion™ High-Fidelity DNA Polymerase (Thermo, F-530XL), and the resultant amplicon was subsequently cloned into pENTR™ 5’-TOPO™ vector (Thermo, K59120). Both clones were confirmed by sequencing before downstream cloning. To generate *UBQ10p:RBP47b::mTb::mVenus* construct (*UBQ10p:RBP47b::PL*), entry vectors carrying the *UBQ10p, RBP47b, pDONR_P2R-P3_R2:mTbID::STOP:L3* (Addgene ID #127356) were recombined into the destination vector pB7m34GW,0 (2). The R4pGWB601 (RIKEN) *UBQ10p:mTbID::NES::mVenus* construct (UBQ10p:PL) used as a control to express cytosolic mTb under the UBQ10 promoter was acquired from Addgene (#127369). Plant expression vectors were confirmed by sequencing and transformed into the Agrobacterium GV3101 strain. To generate *UBQ10p:RBP47b::eYFP* overexpression constructs, entry vectors carrying the UBQ10p and RBP47b sequences were recombined into the R4pGWB640 vector (RIKEN).

### Generation of Arabidopsis transgenic lines

*Arabidopsis thaliana* ecotype Col-0 was used as wild-type (WT). Plant lines for proximity labeling were generated by floral dip ^85^ of Col-0 using the above-described Agrobacterium strains. The selection was done in BASTA (256 μl/L) on soil for T0 offspring. For the next generations, selection on MS plates supplemented with BASTA (10 μg/L) was performed, and transgenic plants were evaluated for fluorescent signal under the confocal microscope. We did not observe any noticeable decrease in viability or developmental delay in our transgenic lines. Based on the combined evaluation of fluorescent signal and gene expression levels, the lines L6 for *UBQ10p:RBP47b::mTb::mVenus* T2, L5 for *UBQ10p::mTb::NES::mVenus*, T2 and L12 for *UBQ10p:RBP47b:eYFP* T3 were selected for further experiments.

### Confocal microscopy

Plants were imaged using a Leica STELLARIS Falcon laser-scanning confocal unit (LEICA Microsystems, Wetzlar, Germany). HC PL APO 63x and 40x 1,2 W CORR UVIS CS2 confocal objectives were used. YFP signal was excited at 515 nm, emission was captured at 525–550 nm, chlorophyll was excited at 650 nm, and emission was captured at 667–755 nm. For Cy5, fluorescence was captured using 651nm (excitation) and 670 (emission). SGs counting was performed on ImageJ using the plugin ComDet v.0.5.5.

### RNA extraction

RNA extraction was performed according to the manufacturer protocol, The Maxwell® RSC Plant DNA Kit (Promega, AS1500). Briefly, 11-day-old seedlings were collected and snap-frozen in liquid nitrogen; frozen tissue was ground in a Retsch Mixer mill until fine powder; 100 mg of ground tissue was mixed with the 600 µL chilled homogenization buffer attached in the Promega kit, including 20 µL/mL 1-thioglycerol. 400 µL of the homogenate solution was transferred into a new tube, and 200 µL Lysis buffer of the Promega kit was added to the sample solution and mixed. The solution was incubated at room temperature for 10 minutes and then centrifuged at 14,000 rpm for 2 min. 500 µL of the supernatant was transferred to the cartridges, and 10 µL of DNase I (attached in the kit) was added to each sample. The samples were set to the Maxwell RSC Instrument (Promega) to start RNA purification. The extracted total RNA was finally dissolved in 50 µL RNase-free water. The extracted RNA concentration was estimated by NanoDrop 2000 (Thermo Fisher Scientific), and the quality was checked by BioAnalyzer 2100 (Agilent Technology, Santa Clara, CA).

### QRT-PCR analysis of gene expression

Total RNA was extracted from seedlings. In brief, 100 mg of ground powder from each tissue was used for RNA isolation using a Maxwell RSC Plant RNA Kit (AS1500) with a Maxwell RSC48 instrument, as indicated above. The RNA yield and quality were assessed using a nanodrop spectrophotometer (Nanodrop Technologies, Wilmington, DE, USA). Quantitative PCR was performed as described in ^86^. Briefly, intron-specific primers (At5g65080) were used to confirm the absence of genomic DNA contamination. First-strand cDNA was synthesized from 2.5 µg total RNA using Superscript III reverse-transcriptase (Invitrogen), and its quality was assessed using primers that amplify 3′ and 5′ regions of GAPDH (At1g13440). qRT-PCR was performed with Quanta Studio 7 (Applied Biosystems, Darmstadt, Germany) in three technical replicates. Reactions contained 2.5 µL 2 × SYBR Green Master Mix (Fast Power SYBR Green; Applied Biosystems), 0.5 µL cDNA (diluted fivefold) and 2 µL of 0.5 µm primers. The sequences of all primers are given in Supporting Information Table S1. To ensure accuracy, primers were first added to each plate, followed by a master mix containing the cDNA, and SYBR Green, RNA, and cDNA quality control reactions were manually pipetted d. Ct values for the genes of interest were normalized by subtracting the mean of four reference genes

### Proximity labelling optimization assay

To confirm the suitability of the proximity labeling assay under heat stress, 11-day-old PL lines, RBP47b^mTb^ and mTb^NES^ were subjected to 22°C, 37°C, 39°C, and 40°C for SGs formation. After selecting 40°C as the working temperature, we tested the effect of 0, 30, 50, and 100 µM of biotin on SGs formation under control conditions and heat stress by submerging the plants in biotin solutions for 1h. Briefly, 15 Plants per treatment were incubated with biotin and washed 2 times with cold water after feeding. Following treatment, the plant material was flash-frozen for later immunoblotting using Streptavidin-HRP according to the manufacturer’s protocol. (Thermo Fisher Scientific). Proteins were extracted as described in ^16^. Coomassie staining was performed as a loading control for SDS gels. Streptavidin immunoblotting revealed only minimal background biotinylation in Col-0 under 30 µM biotin, whereas RBP47bmTb lines displayed a robust labeling profile under biotin application, demonstrating that biotin ligase is fully active under our feeding regimen and functions effectively both at control temperature and during heat stress.

To assess the impact of RBP47b^mTB^ fusion on SG disassembly and set the recovery time point, we performed an SGs disassembly dynamics using confocal microscopy after 1h of heat stress treatment at 40°C supplemented with 30 µM of biotin. The analyzed time points of recovery were as follows: 30, 60, 90, 120, 180 and 240 min. Following treatment, 15 plants per condition were flash-frozen for later immunoblotting using Streptavidin-HRP (Thermo Fisher Scientific). Coomassie staining was performed as a loading control for SDS gels.

### Biotin feeding

11-day-old plants were used for proximity labeling experiments. Biotin feeding was done by submerging the plant material in a 30 μM biotin solution; labeling was performed for 1h for all samples. To elucidate changes in the RBP47b proteome profile during control, heat treatments, and recovery phase, each genotype: Col-0, *UBQ10p:RBP47b::mTb::mVenus*, *UBQ10p::mTb::mVenus* were subjected to biotin feeding at three different time-points: control, heat, and recovery. Biotin feeding for control conditions was performed at e 23°C/light conditions. For the heat treatment, Arabidopsis seedlings were subjected to a pre-heat step of 40°C/dark for 15 min to induce SG formation and avoid biotin depletion by labeling RBP47b proxitome under control conditions. After this step, the biotin feeding was performed, and labeling was continued for 1 h under heat conditions. For the recovery time point, plants were subjected to heat stress at 39°C/dark for 1.15 h and moved to normal growth conditions (light, 120 μM m^−2^ s^−1^ at 23°C). After 60 min of recovery, biotin feeding was performed, and labeling was continued for 1h under recovery conditions.

### Affinity purification of RBP47b proxitome

The affinity purification of biotinylated proteins was performed as described in ^16^ with slight modifications. Briefly, 1 gr per replicate (3 replicates per treatment) of ground material was resuspended in 1 ml of extraction buffer (50 mM Tris pH 7.5, 150 mM NaCl, 0.1% SDS, 1% Triton X-100, 0.5% Na-deoxycholate, 1 mM EGTA, 1 mM DTT, 1x complete, 1 mM PMSF) and incubated at 4°C on a rocking plate for 10 min. 1 μL of Lysonase (Millipore) was added to the suspension and incubated at 4°C for another 15 min. The solution was sonicated 2 times on an ice bath (BRAND) 3 min with 3 min break on ice. The suspension was centrifuged at 15,000 g for 15 min at 4°C to remove the cellular debris. The cleared protein extracts were applied to PD-10 desalting columns according to manufacturer protocols (GE Healthcare) to deplete the free biotin from the samples. The protein concentration was determined by Bradford assay (ThermoFisher) in a 1:10 dilution. Protein concentration was adjusted to ∼8 mg and transferred into a new 2ml low Binding protein tube (Eppendorf), and 100 µL of equilibrated Dynabeads MyOne Streptavidin C1 (Invitrogen) were added to each sample. Halt™ Protease Inhibitor Cocktail (100X) (ThermoFisher) and PMSF were added to a final concentration of 1X and 1 mM, respectively, for overnight incubation at 4°C in a rocking plate platform. The next day, the beads were separated from the protein extract on a magnetic rack, washed, and incubated on the rotor wheel for 4 min with 1 ml of each of the following solutions: 2x with cold extraction buffer (beads were transferred into a new tube the first time), 1x with cold 1 M KCl, 1x with cold 100 mM Na2CO3, 1x with 2M Urea in 10 mM Tris pH 8 at room temperature and with 1 ml 50 mM Tris pH 7.5. Beads were stored at - 80 °C until further processing.

### Sample preparation for Mass Spectrometry

Digestion was performed on the beads. Before digestion, the frozen beads were thawed and washed 2X with 1 mL 50 mM Tris pH 7.5 (transferred to a new tube in the first wash) and 1 ml 2M urea in 50 mM Tris pH 7.5 as described in (3). Beads were resuspended in digestion buffer (6M urea, 2M thiourea in 50 mM Tris pH 7.5) and sonicated 2 times on an ice bath (BRAND) for 3 min with 3 min break on ice. For protein reduction, DTT was added to a final concentration of 10 mM and incubated at 56°C for 30 min with shaking, and then alkylated by adding Iodoacetamide to a final concentration of 20 mM and incubating at 25°C for 30 min with shaking in the dark. After protein denaturation, the urea concentration was lowered to 1 M urea using 50 mM Tris pH 7.5, and a Lys-C/Trypsin mix was added in a 1:25 ratio. The beads were incubated for 10 h at 37°C with shaking. For complete digestion, fresh Lys-C/Trypsin mix was added the following day in a 1:100 ratio and incubated for 3h at 37°C with shaking. The digestion was quenched by adding TFA to a final concentration of 1% and desalted using HyperSep™ C18 Cartridges (Thermo). The C18 columns were washed twice with 1 mL of acetonitrile and then washed twice with 0.1% TFA. The beads and supernatant were loaded into the C18 columns by gravity to allow the binding of the peptides to the matrix. The peptides were washed 3x with 1 mL of 0.1% formic acid. The wash buffer was entirely removed by vacuum prior to elution. The peptides were eluted with an elution buffer (80% acetonitrile, 0.1% formic acid). The desalted peptides were dried in a SpeedVac and stored at -80°C until further processing.

### UPLC-timsTOF Pro 2 QTOF-Based Mass Spectrometry Analysis

Approximately 200 ng of peptide mixture per sample was analyzed using a timsTOF Pro 2 QTOF mass spectrometer coupled with a nanoElute liquid chromatography system (Bruker Daltonik GmbH, Germany). The sample was injected directly into an RP-C18 Aurora emitter column (75 µm i.d.× 250 mm, 1.6 μm, 120 Å pore size) (Ion Opticks, Australia) using a one-column separation method. An 80-min gradient was established using mobile phase A (0.1% formic acid in water) and mobile phase B (0.1% formic acid in Acetonitrile): 2–25% B for 60 min, 25–37% for 10 min, ramping 37% to 95% in 5 min, and maintaining 95% B for 5 min. The column temperature was 50°C, and the flow rate was 250 µl/min. The sample eluting from the separation column was introduced into the mass spectrometer via a CaptiveSpray nano-electrospray ion source (Bruker Daltonik GmbH) with an electrospray voltage of 1.5 kV. The mass spectrometer was set at positive mode with TIMS enabled. The ion source temperature was 180 °C with a dry gas of 3 l/min.

The data acquisition was performed using a PASEF scan as described ^87^. A total of 9 PASEF ramps precursors with a charge between 0-5 with a cycle time of 1.26 s. In each PASEF, the ramp time was set to 120 ms, and the TIMS scan ranged from 0.64 to 1.45 Vs cm^−2^ (1/K0). The collisional energy increased linearly from 20.0 eV at 0.60 (1/K_0_) to 59.00 eV at 1.60 Vs cm^−2^ (1/K_0_). The scan range for both MS and MS/MS spectra was set to 100-1700 *m/z*. The TIMS accumulation time was set to 100 ms. The target precursor intensity is 15,000 au, and the minimum threshold is 1500 au. The active exclusion was for 0.4 min. The mass spectra peak detection uses mass intensity for absolute threshold and mobilogram peak detection with an intensity of 5000.

### Proteome data analysis for Proximity labeling approach and total proteome

To analyze the changes in the proteome of Col-0 and *rbp47abcc’* known contaminants, reversed annotated peptides identified only by one side were removed from further analysis. Proteins with at least two unique peptides were considered for further analysis. LFQ intensities were used in all analyses performed in this study. For group comparison, protein had to be present in at least 3 out of 4 replicates or only under one condition (heat or control). Student T. test (p-value ≤ 0.05) was used to define significantly abundant proteins.

### Protoplast isolation and transformation

Protoplasts were isolated from leaves of 22-day-old Arabidopsis expressing the *UBQ10p:RBP47b:eYFP/Col-0*, as described previously ^88^. The cell walls were digested by incubation in enzymatic solution containing 1% (w/v) Cellulose R-10, 0.25% (w/v) Macerozyme R-10, 20 mM MES-HOK pH 5.7, 400 mM Mannitol, 10 mM CaCl2, 20 mM KCl for 2h. Protoplasts were separated from debris by centrifugation (100 g, 3 min, 4°C); washed two times with ice-cold W5 buffer (154 mM NaCl, 125 mM CaCl2, 5 mM KCl and 2 mM MES-KOH pH 5.7) and resuspended in ice-cold W5 buffer. 100 μl of the protoplast suspension was incubated with PEG 6,000 and 20 µg of the corresponding construct for 15 min at room temperature. Protoplasts were separated from debris by centrifugation (100 g, 3 min, 23°C), resuspended in W5 buffer, and left overnight in the dark at room temperature. The next day, the protoplast was either heated at 42°C for 30 min or kept under room temperature for confocal analysis.

### Sequence analysis

The evolutionary history was inferred by using the Maximum Likelihood method and JTT matrix-based model ^89^. The tree with the highest log likelihood (-2931.59) is shown. Initial tree(s) for the heuristic search were obtained automatically by applying Neighbor-Join and BioNJ algorithms to a matrix of estimated pairwise distances using the JTT model, and then the topology with superior log likelihood value was selected. The tree is drawn to scale, with branch lengths measured in the number of substitutions per site (next to the branches). This analysis involved 4 amino acid sequences. There was a total of 464 positions in the final dataset. Evolutionary analyses were conducted in MEGA11 ^90^ and analyzed using the Jalview java alignment editor ^91^.

### RNA fluorescence in situ hybridization (FISH)

FISH was performed as previously described ^92^. Briefly, 7-day-old plants were under control, and heat stress and stated recovery time points were fixed with 4% paraformaldehyde (PFA) before being squashed. Permeabilization was done in 70% ethanol for 12h. Next, we proceeded with overnight hybridization using Cy5-oligo deoxythymidine (dT) (GeneScript, Taiwan & Middle East Region) at 37 °C. The slides were washed twice before imaging.

### BONCATE

The AHA labeling was performed in 7-day-old roots of seedlings either pre-treated with AHA (HWRK Chem, HWG58500, plants were treated at 42°C for 1h or kept under 23°Ccontrol conditions for recovery plants after heat stress were moved to control conditions for 60 min or 120min. for All samples were immediately frozen in liquid nitrogen after treatments. Samples were pulverized and resuspended in PBS buffer with protease inhibitor cocktail (EDTA free, 1:50 dilution, Roche). After sonication at 4 °C for 40 min, lysates were centrifuged at 18,000g for 10 min to collect the supernatant and quantified by the Bradford assay. For click reactions, 20 μg protein was incubated with 60 μl PBS buffer containing 100 μM biotin-alkyne (Sigma, 764213-5MG), 333 μM CuSO4, 666 μM Tris(3-hydroxypropyltriazolylmethyl) amine (Sigma, 762342-100MG), and 0.5 mg ml−1 sodium ascorbate in the dark for 1.5 h. Then, the reaction was stopped by adding PBS, methanol, chloroform, and water step by step. After centrifugation at 14,000g for 2 min, the protein layer was collected and precipitated using methanol. After centrifugation at 18,000g for 10 min, the pellet was collected, washed twice with methanol, dried at room temperature and dissolved using 2 M urea by sonication at 50 °C. After quantifying the protein concentration by the Bradford assay, 0.25 μg protein per 50 μl PBS was coated in a 96-well high-binding plate (Corning) per well at 4 °C overnight for ELISA. Unbound protein was washed out using PBS with Tween 20 (PBST) buffer, and a blocking solution (2% BSA dissolved in PBST) was added per well. After incubation at 37 °C for 1 h, the blocking buffer was discarded, and streptavidin–horseradish peroxidase (HRP) (Abcam, 1:60,000 diluted in PBST) was added. After incubation at room temperature for 40 min, antibodies were discarded, and the well was washed five times with PBST. Antibody binding was detected by the addition of HRP substrate (0.1% o-phenylenediamine, 0.075% H2O2, 0.1 M citric acid and 0.2 M Na2HPO4, pH 5.6) per well and incubation in the dark at room temperature for 8 min. The addition of 2 M H2SO4 stopped the reaction, and the absorbance at 450 nm was detected by a microplate reader (BioTek Cytation5).

### Phenotyping and Heat Stress Treatment

Seeds for Arabidopsis (*Arabidopsis thaliana*) ecotype Columbia-0 (Col-0), *hsp101* (AT1G74310; hot 1-3; NASC ID: N16284), and the homozygous CRISPR (Cas)9-free *rbp47abcc′* line were surface sterilized and grown in plates as described in the Plant Material and Growth Conditions section. Control plates were kept in an LED Plant Growth Chamber (Percival Scientific, Inc., Perry, Iowa) at 23 °C in long-day conditions (16h light/8h dark with 120 μM m−2 s−1 for the whole experiment. For heat shock, 11-day-old plants were exposed to 42 °C for 5 h in a preheated oven to observe heat shock response (80 plants in total). After the exposure to stress, the plants were placed back in the growth chamber at normal conditions (23 °C in long-day conditions (16h light/8h dark with 120 μM m^−2^ s^−1^) for 7 days to recover after stress. Plates were photographed using a Canon model EOS 90D (www.canon.com) fitted with a Canon EFS 18-135mm lens.

### Total Chlorophyll content

The control and treated genotypes underwent fresh weight measurement and aerial part collection for total chlorophyll content analysis. Each of the wells in 6-well sterile polystyrene culture plates was filled with an equal amount of freshly made 80% ethanol. The aerial parts of each genotype were entirely submerged in a separate well and kept overnight in the dark with gentle shaking. The absorbance of the extracted solution was determined spectrophotometrically using the plate reader THERMO Varioskan Flash spectrophotometer set to measure values at wavelengths of A664 and A647 as described by ^93^. The concentration of total chlorophyll content was calculated as ((7.93A_664_) + (19.53A_647_))/fresh weight (μmol/g). Reference blanks were used for calculation.

### High-Throughput phenotyping

The Arabidopsis (*Arabidopsis thaliana*) ecotype Columbia-0 (Col-0), *hsp101* (AT1G74310; *hot1-3*; NASC ID: N16284), and the homozygous CRISPR (Cas)9-free *rbp47abcc′* t were sterilized and grown in plates as described previously. Peat soil (Stender, Germany) was mixed with perlite and vermiculite (ratio 3:1:1), double autoclaved, and filled in PSI-compatible pots (Photon Systems Instruments, Czech Republic) to 100g (±1.0g). The pots were watered to full saturation level with tap water and placed overnight to drain excess water. When the pots were at the proper moisture level, 10-day-old seedlings of uniform size were individually transplanted into prepared pots. A blue mat was placed on top of the soil to avoid background reflection of the soil that could influence proper chlorophyll segmentation measurements. The pots were placed in an environmentally controlled PSI-growth room and adapted to humidity by covering the trays with an individually transparent plastic lid cover for 3 days after the lid was removed. The PSI growth room maintained a constant temperature of 22°C, with a sensor sensitivity range of ±0.1 °C, and sustained a relative humidity of 60%, with a sensor sensitivity range of ±1%, and a carbon dioxide concentration of 400 ppm, with a sensor sensitivity range of ±100 ppm. The plants were cultivated in a 16 h/8 h light/dark cycle with a 120 µmol/m2 light intensity. A total of 152 plants were used for the phenotyping, including 51 Col-0, 54 *rbp47abcc’*, and 47 *hsp101* genotypes. Each PSI tray (5×4 pot space per trey) was filled with a total of 10 plants in a random distribution and introduced into the data bank of the PlantScreen^TM^ Compact System (PSI, Czech Republic). Each PSI tray (configured with a 5 × 4 pot layout) contained a total of 10 plants, randomly distributed across the available spaces. The trays were housed inside a completely automated PSI growth chamber, an enclosed system with conveyor belts, integrated imaging RGB cameras, weighing platforms, and a watering station for high-throughput phenotyping. These trays were integrated into the PlantScreen™ Compact System (PSI, Czech Republic), an automated system that maintained optimal soil-water levels by adjusting each pot’s moisture to a pre-set reference weight. Upon placement in the PSI growth chamber, the system continuously conducted daily weighing and watering cycles to ensure consistent hydration. Hence, they underwent automated cycles of weighing and watering daily to reach the soil-water reference weight. For phenotyping, two days before heat treatment, the PlantScreen^TM^ system was set to take RGB images and measurements for all analyzed parameters at three periods during the day (12 AM, 8 AM, and 6 PM). During the heat treatment process, when the plant varieties reached 22 day old, half the total number of trays were relocated to an environmentally controlled Conviron growth chamber (Conviron GEN2000 TA, Conviron, Canada), preheated to 45°C overnight. After a 9h stress treatment, the pots were reinstated in the PSI growth room for subsequent plant phenotyping analysis. The phenotyping results were collected and analyzed with R package to assess the genotype’s responses to heat stress.

### RNA-seq analysis

Library sequencing was performed by Novogene, and the resultant FASTQ files were used for analysis via a standard RNA-seq pipeline with slight modification ^94^. Briefly, fastp was used to trim raw reads and filter out those of low quality ^95^. Trimmed reads were then mapped to the Arabidopsis genome ^96^ using HISAT2 ^97^, and bam files were generated using samtools^98^. Mapped reads were then assigned to gene features in the Araport11 genome annotation using featureCounts ^99^, and the resulting reads counts file was used as input for differential expression analysis. DESeq2 v1.46.0 ^100^ was used to identify differentially abundant genes in *rbp47abcc’* vs. Col-0 comparisons in each condition and heat vs. control and recovery vs. heat comparisons for each genotype. GO Enrichment analysis was performed using the clusterProfiler v4.14.6 ^101^ R package and GO annotations from the Bioconductor org.At.tair.db (v3.20.0) annotation package.

### Methyl Tert-Butyl Ether -Methanol (3:1) Extraction Method and Sample Preparation

We follow the protocol described by ^102^ with slight modifications for protein, metabolites, and lipid extraction. 11-day-old plants were used for this experiment. Briefly, ∼160 mg of homogenized tissue was resuspended in one of the pre-cooled (-20 °C) extraction solvent mixture 1 (M1), which consisted of 75 ml of methyl tert-butyl ether and 25 ml of methanol (3:1, vol/vol). After adding the M1 extraction solvent, the tubes were thoroughly vortexed for 1 min and then incubated on an orbital shaker (100 rpm) for 45 min at 4 °C, followed by a 15-minute sonication step in an ice-cold bath. For phase separation, 650 µl of solvent M2 (75 ml of water, 25 ml of methanol, 3:1, vol/vol) was added to each vial/tube, and the samples were again thoroughly vortexed for 1 min. After that, the samples are centrifuged at 20,000g for 5 min at 4 °C. Three phases were observed at this point: the upper phase (500 µL) corresponds to the lipid-containing phase, and ∼600 µL of the lower phase corresponds to the metabolites-containing phase. Each phase was transferred into a new pre-labeled 1.5 mL tube and was evaporated using a SpeedVac concentrator at RT. After removing both phases, the remaining protein/starch pellet was washed by adding 500 µl methanol and thoroughly vortexing the samples for 30 s. The samples were centrifuged at 20,000g for 5 min at 4 °C. This washing step was repeated two more times. All dry pellets from fractions and proteins were kept at -80 °C until further processing.

### Proteome extraction

For protein extraction, the pellet was resuspended in 200 µl of 8M urea, 2 M thiourea in 50 mM of ammonium bicarbonate supplemented with 0.03% of ProteaseMAX™ Surfactant, 1x cOmplete™, Mini, EDTA-free Protease Inhibitor Cocktail. Samples were incubated for 60 min onto an orbital shaker set at 300 rpm. The samples were centrifuged at 10,000 g for 10 min to remove further cell debris. The supernatant was transferred into a new low-protein binding tube of 1.5 mL; the proteins were denatured by incubating them with 10 mM DTT at 56 °C, 600 rpm for 30 min, followed by the addition of 20 mM Iodoacetamide, and samples were kept in the dark at room temperature and shaking, 600 rpm. Samples were diluted with 50 mM of ammonium bicarbonate to a final concentration of < 1 M urea. Protein concentration was determined using the Bradford assay (Reference). According to the manufacturer’s protocol, 150 µg of protein extract were digested in-solution using a Trypsin/Lys-C mixture (Mass Spec Grade, Promega) in a 1:25 ratio. After the digestion, the samples were desalted using C18 stage tips as described in ^103^. After the elution of the digested and desalted peptides from C18-stage tips, the samples were concentrated to near dryness in a SpeedVac; dry peptides were stored at -80 °C until further processing. Peptides were resuspended in 0.1% formic acid, and concentration was adjusted to 500 ng/µL; the peptide mixtures were analyzed by LC-MS/MS using LC-MS.

### LC-MS for Secondary Metabolomics and data analysis

400 µl of the polar phase was transferred into a pre-labeled 1.5 ml microcentrifuge tube, and the samples were dried down in a SpeedVac concentrator without heating. For the direct analysis, the samples were handled as described previously in Giavalisco et al., 2023. Briefly, the dried pellets of the polar phase were re-suspended in 150 µL UPLC-grade methanol: water (8:2, vol/vol); the resuspended samples were centrifuged at 16,000 g 4 °C for 5 min and transferred to labeled HPLC vials with glass insert. For LC-MS, untargeted metabolomics samples were analyzed using a Vanquish Ultra-High Performance Liquid Chromatography (UHPLC) system (Thermo Scientific™) coupled to an Orbitrap ID-X mass spectrometer (Thermo, Germany) at the Analytical Chemistry Core Lab (KAUST, Saudi Arabia). Sample extracts were separated on an HSS T3 UPLC column (150 x 2.1 mm, 1.7 µm, Thermo Scientific) using a mobile phase consisting of (A) H2O 0.1% formic acid and (B) Acetonitrile 0.1% formic acid.

The gradient used for the analysis is as follows: 0 – 1.5 min, 0%B, 1.5 – 15 min, 0 to 100% B, 15 – 17 min, 100%B, 17 – 17.5 min, 100% to 0% B, 17.5 – 21 min, 1%B. The injection volume for all samples was 10 µL, and the flow rate was set at 0.400 mL/min. The column oven and autosampler temperatures were held at 35°C and 5°C, respectively. The mass spectrometer was operated in positive and negative polarity switching mode. The spray voltage of the heated electrospray (HESI) source was set at 3500 V and 2500 V for positive and negative modes, respectively. The vaporizer temperature was maintained at 350°C, and the ion transfer tube was set to 300°C. The sheath gas was set to 50 (arb), auxiliary gas to 10 (arb), and sweep gas to 1 (arb).

The acquisition was performed in a mass range of 100 to 1000 m/z using the AcquireX data-dependent acquisition (DDA). The full scan resolution was set to 120000 full-width half maximum (FWHM), at *m/z* 200, and MS/MS was acquired at 30000 resolution FWHM with a stepped collision energy of 10%, 30%, and 50% normalized collision energy (NCE). The acquired raw data were processed using Compound Discoverer software (version 3.3, Thermo Scientific). For comprehensive metabolomics data analysis we used MetaboAnalyst 6.0. ^52^, and identification was performed using online and in-house mass spectral libraries and databases.

### GC-MS for Primary Metabolomics and data Analysis

A Thermo Scientific Orbitrap Exploris GC 240, with a maximum resolving power of 240,000 at *m/z* 200 FWHM, was utilized for primary metabolite analysis. Sample introduction was conducted using a Thermo Scientific™ TriPlus™ RSH auto-sampler. Chromatographic separation was achieved with a Thermo Scientific™ TRACE™ 1310 Gas Chromatograph, equipped with a Thermo Scientific™ Trace GOLD™ TG-5SilMS (30 m × 0.25 mm i.d. × 0.25 μm) column. The instrument was calibrated with PFTBA to ensure a mass accuracy of <1.0 ppm.

The mass spectrometer operated in electron ionization (EI) mode, with an ion source temperature of 300 °C and electron energy of 70 eV at an emission current of 50 µA. The full scan acquisition mode spanned a range of 35-700 Da, with an orbitrap resolution of 60,000 FWHM (measured at m/z 200). The automatic gain control (AGC) target was set to standard, and the maximum injection time was in auto mode. Data acquisition employed lock-mass correction with the GC column bleed siloxane mass at *m/z* 207.

For sample injection, a volume of 1 µL was introduced using a 10 µL syringe into a Thermo Scientific™ Liner GOLD single gooseneck with glass wool, under a split flow of 40 mL/min and a purge flow of 5 mL/min. The helium carrier gas was maintained at 1.2 mL/min. The oven temperature started at 70 °C (held for 2 minutes), ramped to 220 °C at 8 °C/min, and then increased to 325 °C at 16 °C/min, where it was held for 10 minutes.

### Sample Preparation for GS-MS analysis

Sample preparation followed an adapted protocol from ^102^ Samples were derivatized with BSTFA (O-bis(trimethyl-silyl)trifluoroacetamide, Sigma-Aldrich, USA), spiked with a 10 µg/mL stock solution of C7-C40 hydrocarbons (Supelco, 49452-U). A pooled quality control sample was prepared by combining aliquots from all extracted samples. Sample vials were analyzed in a randomized order to minimize result bias.

### Statistical and Multivariate Analysis of GS-MS profile

The raw data were processed using Thermo Scientific Compound Discoverer 3.3 to perform chemometric analyses, including spectral deconvolution, compound identification, and multivariate statistical evaluation. For comprehensive metabolomics data analysis, we used MetaboAnalyst 6.0 ^52^. An untargeted metabolomics workflow was applied to generate principal component analysis (PCA) and volcano plots, enabling the identification of significant features based on their m/z values and retention times. Peak data were deconvoluted, aligned, filtered, and annotated through spectral library matching against NIST20 and the Orbitrap GC-MS HRAM Metabolomics Library.

### LC-MS for Lipidomics

Samples were analyzed using a Vanquish Ultra-High Performance Liquid Chromatography (UHPLC) system (Thermo Scientific™) coupled to a timsTOF Pro 2 mass spectrometer (Bruker Daltonics, Germany). Sample extracts were separated on an accucore C30 UHPLC column (150 x 2.1 mm, 2.6 µm, Thermo Scientific) using a mobile phase consisting of (A) acetonitrile/water (60:40), 10 mM ammonium formate, 0.1% formic acid and (B) isopropanol/acetonitrile (90:10), 10 mM ammonium formate, 0.1% formic acid. H2O with 0.1% formic acid (A) and acetonitrile with 0.1% formic acid (B). The gradient used for the analysis is as follows: 0 – 2 min, 30 to 43%B, 2 – 2.1 min, 43 to 55% B, 2.1 – 12 min, 55 to 65%B, 12 – 18 min, 65 to 85% B, 18 – 20 min, 85 to 100%B, 20 – 25 min, 100%B, 25 – 25.1 min, 100 to 30%B, 25.1 – 31 min, 30%B. The injection volume for all samples was 10 µL, and the flow rate was set at 0.260 mL/min. The column oven and autosampler temperatures were held at 45°C and 5°C, respectively.

The mass spectrometer was operated in positive and negative modes using a vacuum-insulated probe – heated electrospray (VIP-HESI) source. The capillary voltage was set at 4500 V for positive and negative, the nebulizer gas pressure was set to 3.5 Bar, and the dry gas and sheath gas flow were set to 10 L/min and 5.0 L/min, respectively. The dry temperature and sheath gas temperature were maintained at 260°C and 350°C, respectively. The acquisition was performed in a mass range of 20 to 1200 m/z using Parallel Accumulation-Serial Fragmentation (PASEF) and the ion mobility range was set from 0.55 to 1.86 1/K0. The MS/MS was acquired with a stepped collision energy of 20 eV to 40 eV. The acquired raw data were processed using Metaboscape software version 2023b (Bruker Daltonics, Germany). For comprehensive metabolomics data analysis, we used MetaboAnalyst 6.0. ^52^

### Reactive Oxygen Species detection assay

Staining with 5-(and 6)-carboxy-2′,7′-dichloro dihydrofluorescein diacetate (DCFH-DA), 3,3′-diaminobenzidine (DAB) was used to determine ROS accumulation. In all cases, 15 plants per tested line were used, and each line was evaluated under control, heat, and 5 μM MeJA. For the determination of ROS production with DCFH-DA staining done as described previously ^104^, 7-day-old plants were immersed in 60 µM DCF-DA in a standard medium (1 mM KCl, 1 mM MgCl2, 1 mM CaCl2, 5 mM 2 mM MES 2-(N-morpholino) ethanesulfonic acid pH 6.1). The plants were then washed 2x and observed under a GFP filter microscope (excitation 480 nm, emission 527 nm). Fluorescent quantification was performed using ImageJ. H_2_O_2_ accumulation was detected using DAB (3,3′-diaminobenzidine) staining as described before ^105^. Briefly, plants were immersed in a solution of DAB (1 mg/mL) and placed in the dark for 4 h. After this time; the solution was removed, and a solution of ethanol: acetic acid: glycerol (3:1:1) was added for 12 h. The presence of a brown precipitate, indicating H_2_O_2_ production in the leaves, was then observed for qualitative determination.

### Statistical analysis

Statistical analyses and graphs were performed using GraphPad software ^106^ or R-

### Data availability

Proteomic Data were deposited to the ProteomeXchange Consortium via the PRIDE partner repository ^107,108^ with the dataset identifier: 1-20250515-085048-3250000

## Acknowledgments

We thank Dalila Bensaddek and Huoming Zhang from the KAUST proteomic core lab for their help with sample injection and discussions for proximity labeling optimization. We thank Klara Panzarowa and the PSI team for helping with the experimental design and remote monitoring of the facility. We are grateful to KAUST Plant Growth Core Lab Facility members, especially John Rahmer and Angelo Gallone, for their technical assistance during PSI experiments.

## Author Contributions

M.C. conceptualized the project, supervised the project, advised the experimental strategy, analyzed the data, and wrote the manuscript; I.E.H.S. devised the experimental strategy, optimized the proximity labeling technique, performed confocal microscopy, performed omics experiment and prepared samples for RNAseq, proteomic and metabolomics, analyzed the data, wrote manuscript and visualized the results; I.T. performed phenotyping experiments including high-throughput phenotyping with PSI, analyzed the data, assisted with writing the manuscript and visualization; A.K., performed RNAseq analysis and visualization, assisted with writing the manuscript and visualization; A.D.L.N. and A.C.N.D. assisted with RNAseq data analysis, visualization and fruitful discussions; F.M.S. performed ribosome subunit mapping and analyzed proteomic data, data visualization; O.S.R. supervised F.M.S. analysis; I.M.L. assisted with proximity labeling and omics experiment, assisted with review and editing of the manuscript; W.W. and Y.C. performed BONCATE experiment; A.S. prepared and run metabolomics samples (LC-MS), assisted with data analysis and discussion; S.S. prepared and run metabolomics samples (GC-MS), assisted with data analysis and discussion; All authors contributed to the final manuscript preparation.

