## Supplementary figures for "RBP47 family members are negative regulators of heat stress tolerance in *Arabidopsis thaliana*"

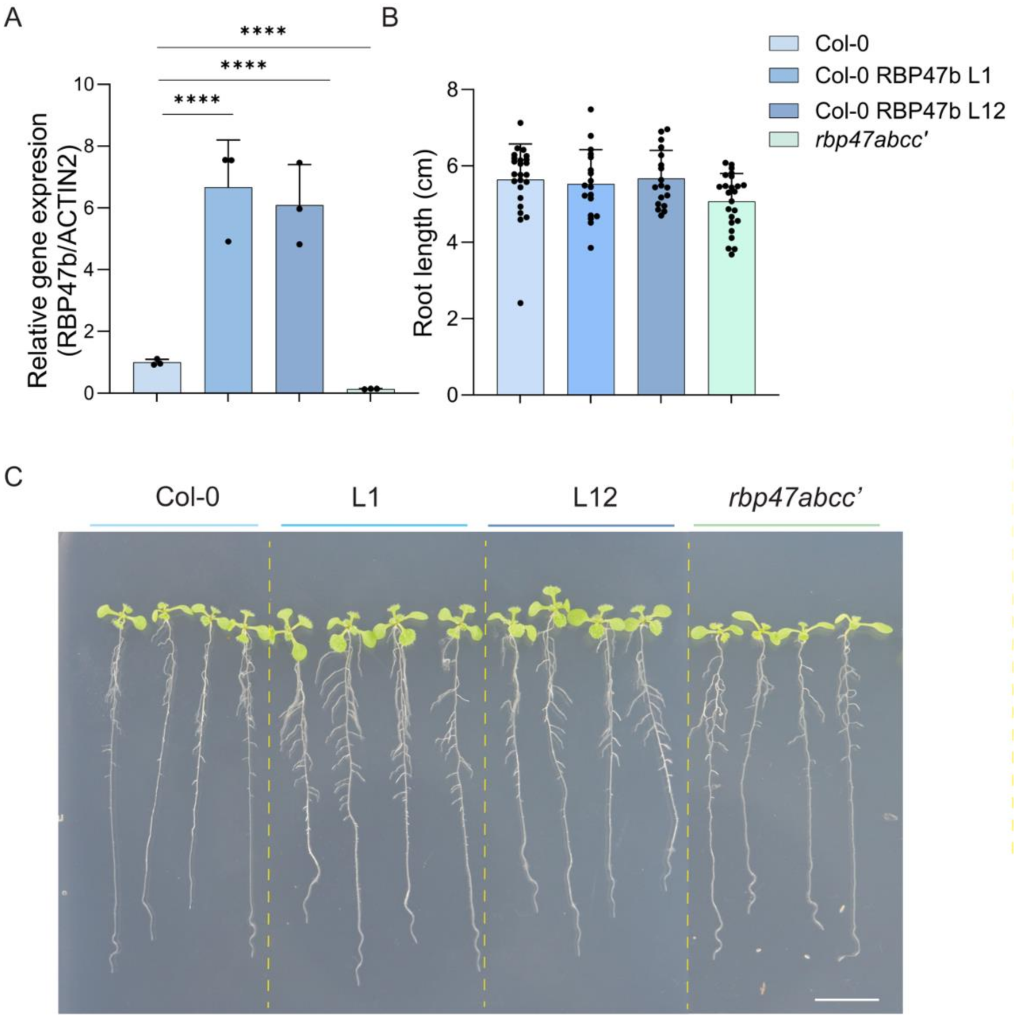


**Supplementary Figure 1. Phenotypical characterization and gene expression analysis of *RBP47b* overexpression lines (in the Col-0 background) and *rbp47abcc'* mutant plants.** **A**. Quantification of *RBP47b* transcript levels in 11-day-old Col-0, *UBQ10p::RBP47B-YFP* (L1 and L12), and *rbp47abcc'* plants. Transcript levels were normalized to *ACT2*, and error bars represent the mean ± SD of triplicate reactions. **B**. Primary root length (cm) of the genotypes from panel **A**. Bar plots show mean ± SD ($n$ = 20, from three independent experiments). Statistical significance was assessed using one-way ANOVA followed by Dunnett's multiple comparisons test ($p < 0.05$; $p < 0.01$; *$p < 0.001$; **$p < 0.0001$). **C**. Representative images of 11-day-old Col-0, *pUBQ10::RBP47B::YFP*, and *rbp47abcc'* mutant plants. Scale bar, 1 cm.

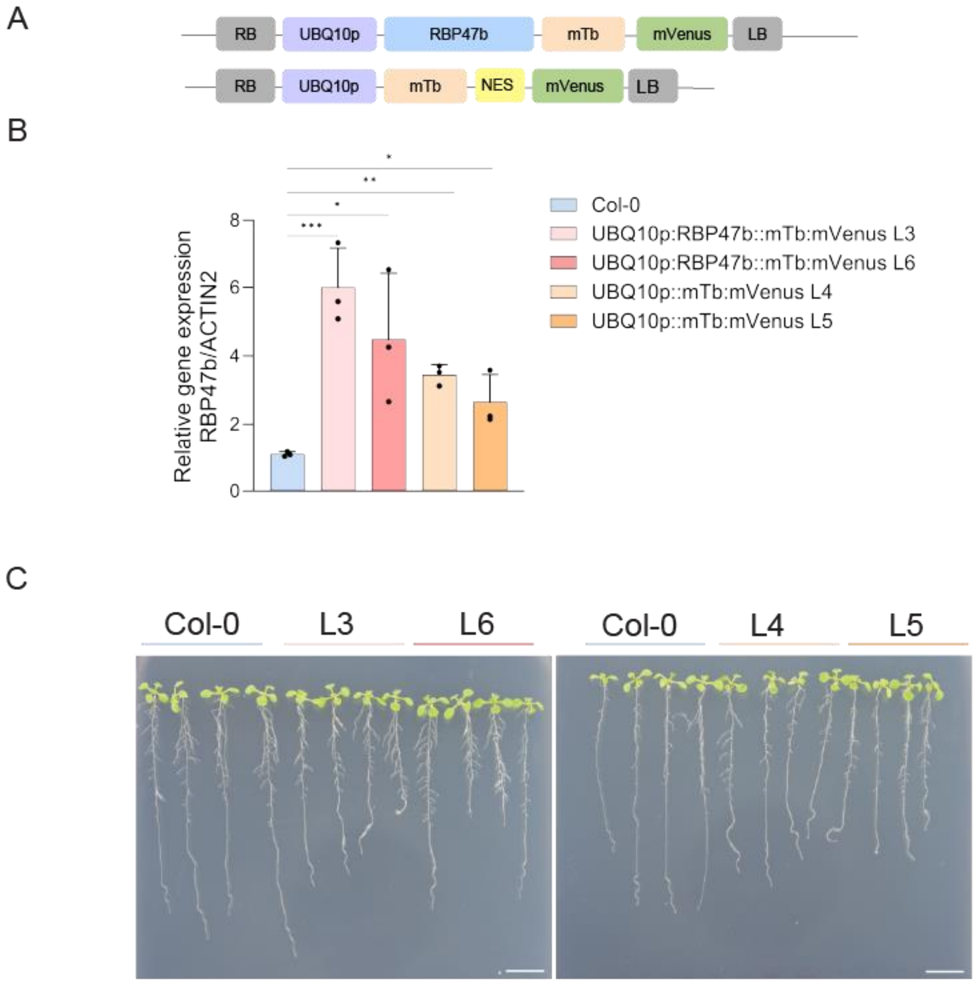


**Supplementary Figure 2. Gene expression profile and phenotype analysis of *UBQ10p:RBP47b::mTb::mVenus* and *UBQ10p::mTb::NES::mVenus* lines**. **A.** Schematic representation of the *UBQ10p:RBP47b::mTb::mVenus* construct. **B**. Relative RBP47B transcript levels in 11-day-old Col-0 seedlings and transgenic lines expressing *UBQ10p:RBP47b::mTb::mVenus* (L3, L6) and *UBQ10p::mTb::NES::mVenus* (L4, L5). Transcript levels were normalized to the expression of the housekeeping gene *ACT2*. Bar plots show mean ± SD (n = 3, from three biological replicates). **C**. Representative images of proximity labeling lines. Scale bar, 1 cm. Bar plots represent mean ± SD (*n* = 5). Statistical significance in **A** was determined by one-way ANOVA followed by Dunnett's multiple comparisons test (*p < 0.05; **p < 0.01; ***p < 0.001).

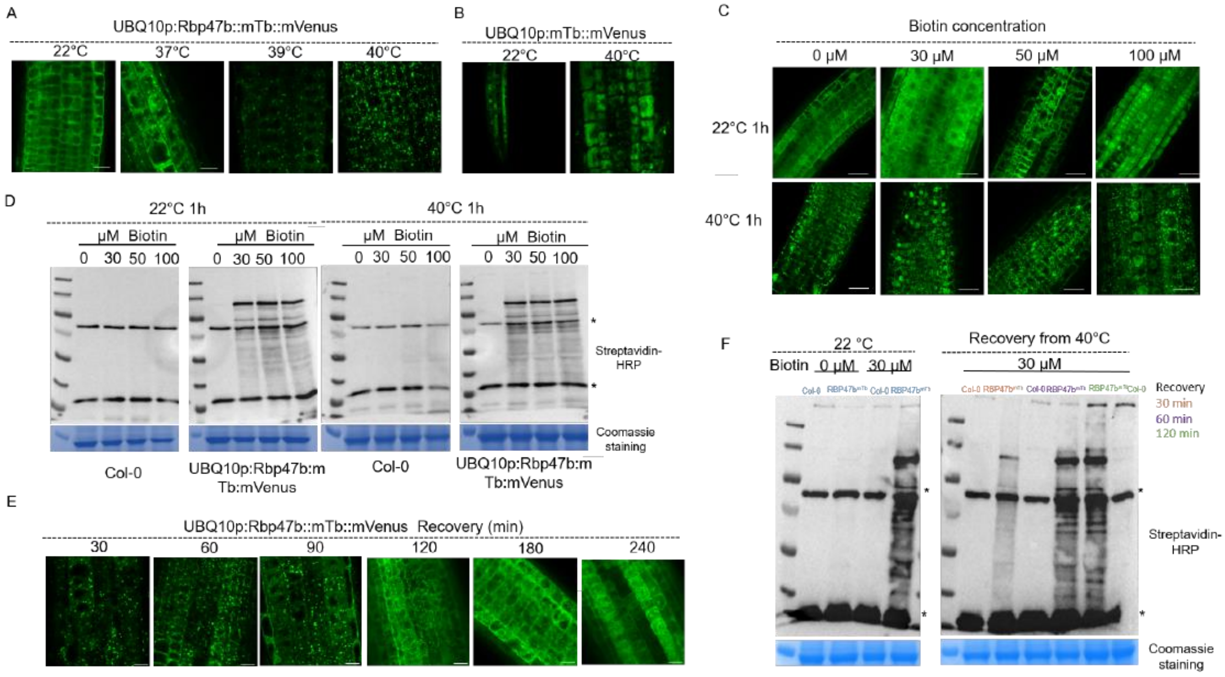


**Supplementary Figure 3. Biotinylation profiles of UBQ10p:RBP47b::mTb::mVenus (L6) and *UBQ10p::mTb::NES::mVenus* (L5) transgenic lines under different concentrations of biotin for the proximity labeling assay under control, heat, and recovery. A**. Fluorescence imaging of *UBQ10p:RBP47b::mTb::mVenus* root cells from 5-day-old seedlings under control conditions (22°C) and heat stress (37°C, 39°C, and 40°C) for 1h. **B**. Fluorescence imaging of *UBQ10p:mTb::NES::mVenus* root cells from 5-day-old seedlings under control conditions (22°C) and heat stress (40°C) for 1h. **C**. Fluorescence imaging of *UBQ10p:RBP47b::mTb:mVenus* root cells from 5-day-old seedlings treated with increasing Biotin concentrations (0, 30, 50, 100 μM) under control (22°C) and heat stress (40°C for 1h) conditions. **D**. Western blot analysis showing biotinylation profiles in Arabidopsis Col-0 plants and *UBQ10p:RBP47b::mTb::mVenus* seedlings treated with increasing biotin concentrations used in (C). Biotinylated proteins were detected using streptavidin-HRP. Coomassie Brilliant Blue staining is shown as a loading control. Asterisks indicate naturally biotinylated proteins. **E**. Confocal imaging showing the *UBQ10p:RBP47b::mTb::mVenus* disassemble dynamics from 5-day-old roots after heat stress (40°C for 1h). **F**. Western blot analysis of biotinylation profile in 5-day-old seedlings incubated at 22°C for 1h. Biotinylated proteins were detected using streptavidin-HRP. Coomassie Brilliant Blue staining is shown as a loading control. Asterisks indicate naturally biotinylated proteins.

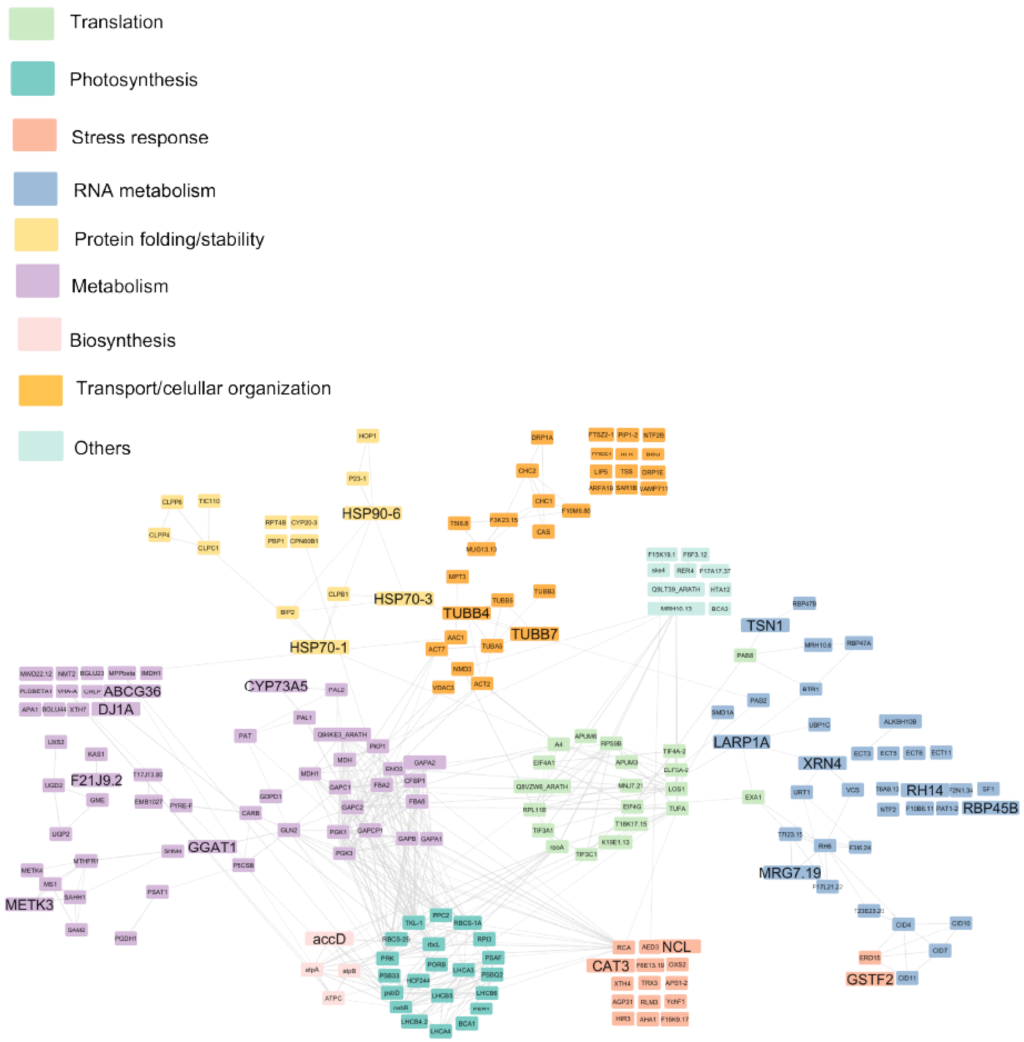


**Supplementary Figure 4. RBP47b protein proxytome network under control conditions identified through proximity labeling.** Network representation of significantly enriched RBP47b interactors (p-value ≤ 0.05, fold change ≥ 1.5) detected under control conditions. Nodes represent proteins associated with RBP47b, and edges denote their interactions. Functional categories are color-coded according to enriched Gene Ontology (GO) terms, with interactors grouped based on their biological roles. Categories include translation (green), photosynthesis (cyan), stress response (orange), RNA metabolism (blue), protein folding/stability (purple), metabolism (lavender), biosynthesis (yellow), transport/cellular organization (dark orange), and others (light cyan). Network visualization was performed using Cytoscape software.

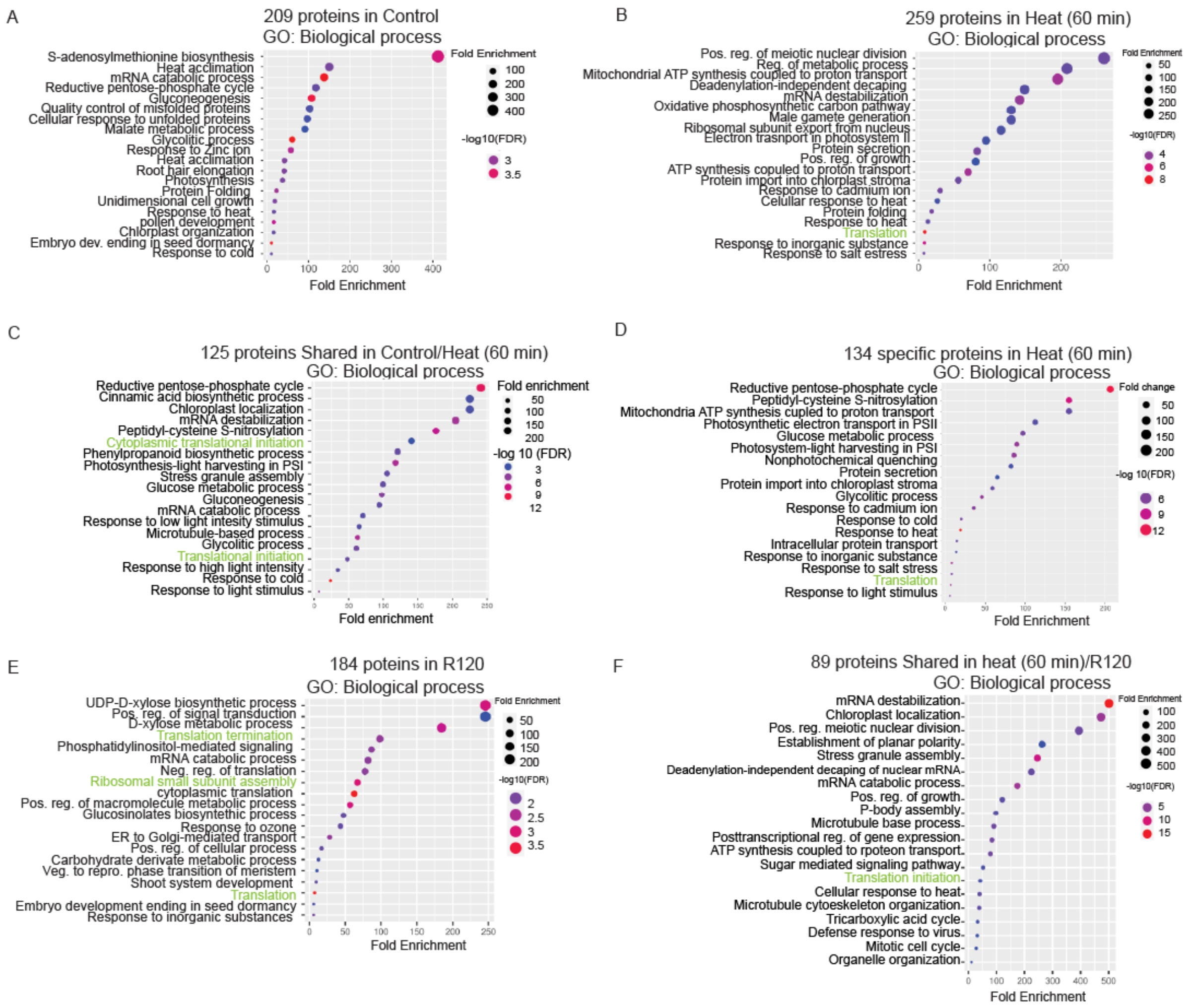


**Supplementary Figure 5. Gene Ontology (GO) enrichment analysis of shared and unique biological processes in the RBP47b protein proxytome.** The top 20 enriched biological process GO terms were obtained with ShinyGo 0.82 software of the RBP47b protein interactome, which was identified by proximity labeling in 11-day-old seedlings. **A**. under control conditions. **B**. Heat stress. **C**. Shared between control and heat **D**. Heat specific **E**. Recovery for 120 minutes (R120). **F**. shared between heat 60 min (specific) and R120. Dot size represents fold enrichment, while color intensity indicates statistical significance (-log10 FDR). Translational processes are highlighted in green.

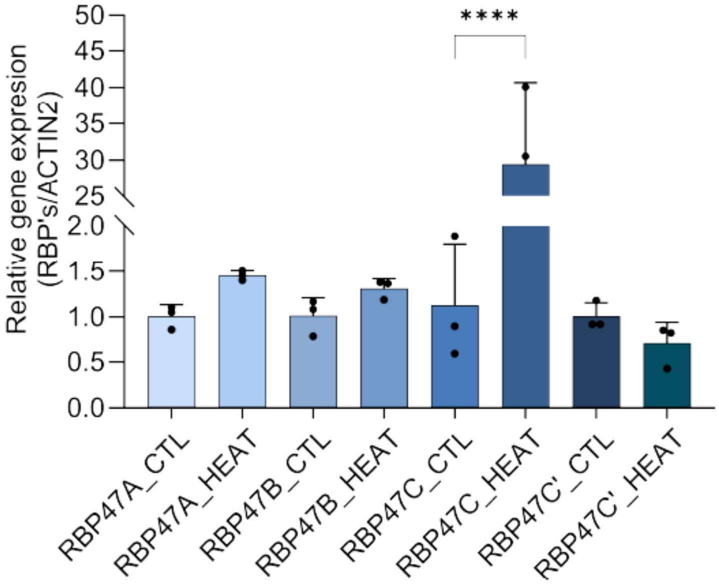


**Supplementary Figure 6. Gene expression analysis of the RBP47 family members suggests a functional redundancy.** Total RNA extracted from 11-day-old Col-0 under control and heat conditions. Data analysis was performed using the delta CT (critical threshold) method, with ACT2 mRNA levels as the internal reference. Three biological replicates (n=3) were included in this analysis. Data represents the mean ± SD (n = 9, from three independent experiments). Statistical significance was assessed using one-way ANOVA followed by Dunnett's multiple comparisons test ($p < 0.05$; $p < 0.01$; *$p < 0.001$; **$p < 0.0001$).

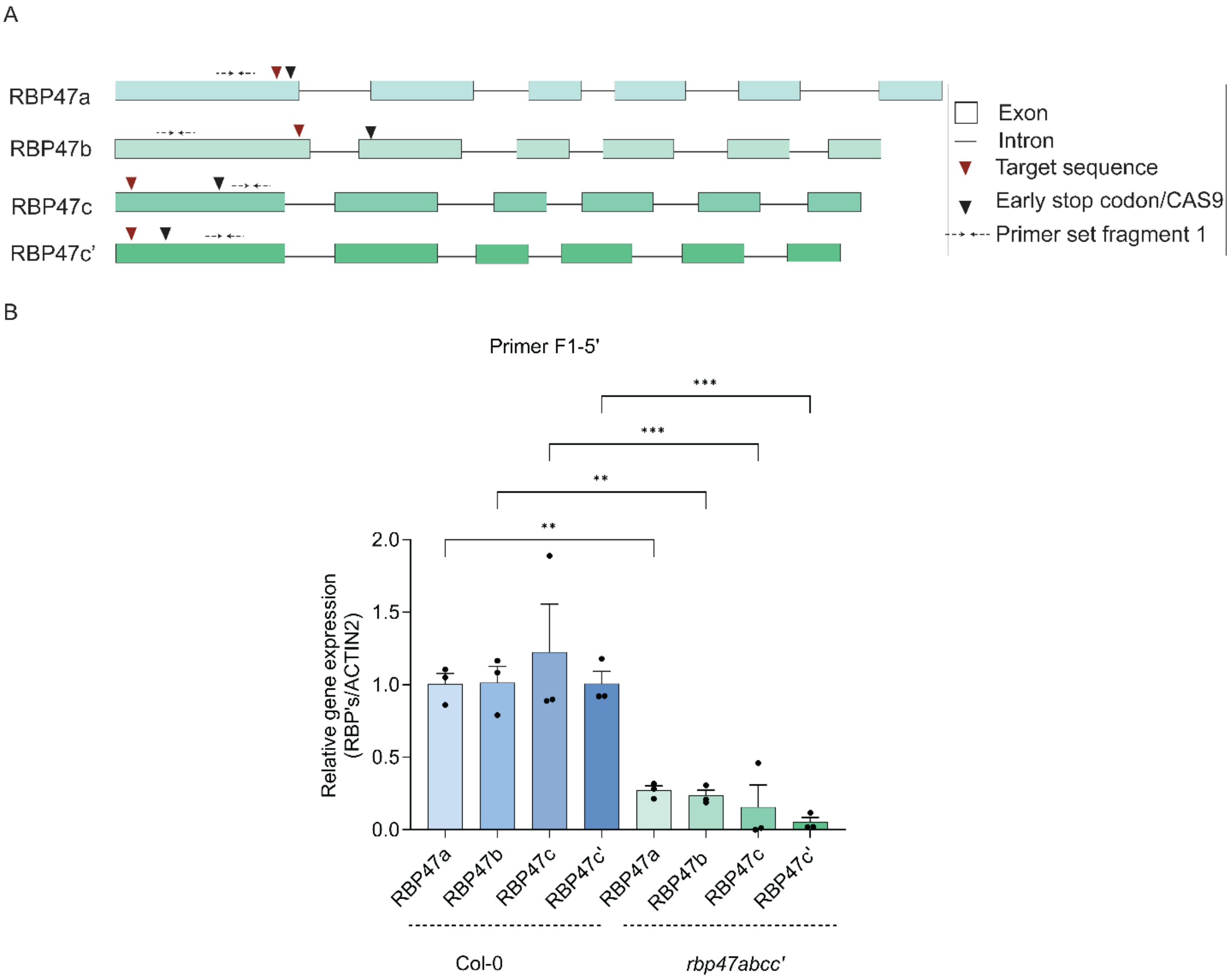


**Supplementary Figure 7. Gene expression analysis of RBP47 family members in Col-0 and *rbp47abcc mutant* backgrounds. A**. Schematic representation of the CRISPR/Cas9 target sites *in rbp47a, rbp47b, rbp47c, and rbp47c'* genes rectangles represent exons, lines correspond to introns, red triangles represent target sequences for Cas9, black triangles indicate where the early stop codon was generated, and sashed lines indicates the binding site of primers used for gene expression. **B**. Relative gene expression levels of *RBP47a, RBP47b, RBP47c,* and *RBP47c'* in Col-0 and *rbp47abcc'* mutant 11-day-old seedlings, as determined by qRT-PCR. *ACTIN2*, a constitutively expressed gene, was used as an internal control. Data represents the mean ± SD (n = 9, from three independent experiments). Statistical significance was assessed using one-way ANOVA followed by Dunnett's multiple comparisons test ($p < 0.05$; $p < 0.01$; *$p < 0.001$; **$p < 0.0001$).

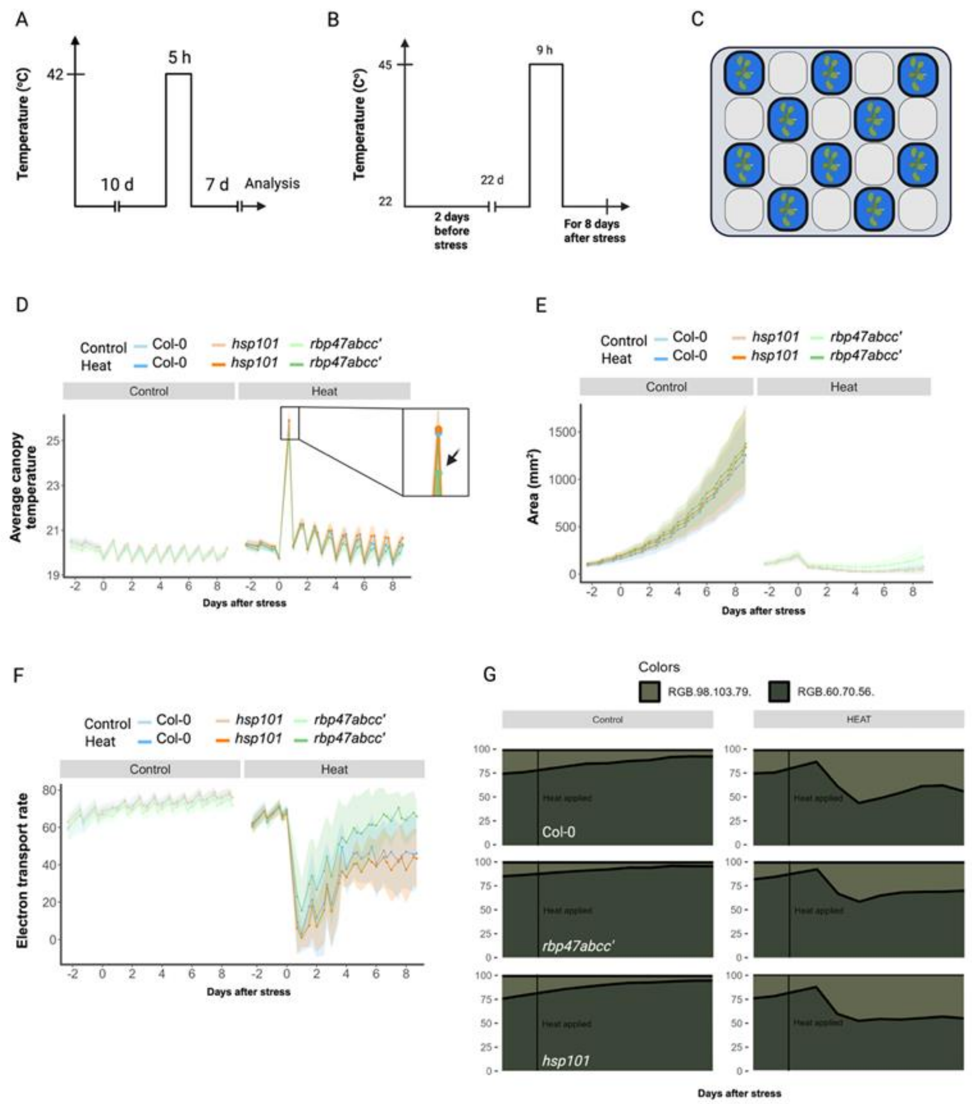


**Supplementary Figure 8. Sustained growth and photosynthetic function in *rbp47abcc'* during recovery from heat stress. A.** Schematic design of heat stress conditions applied to 11-day-old seedlings. **B.** Schematic design of heat stress conditions applied to 22-day-old plants using a high-throughput phenotyping approach via PSI. **C.** Tray layout indicating the spatial distribution of Col-0, *hsp101*, and *rbp47abcc'* genotypes during high-throughput phenotyping. **D.** Average canopy temperature of the 22-day-old plants (Col-0, *hsp101*, and *rbp47abcc'*) analyzed by PSI during heat stress. **E.** Measurement of rosette area ($mm^2$) used as a proxy for growth of the 22-day-old heat-treated plants analyzed via PSI. **F.** Electron transport rate (ETR) as a measure of photosynthetic activity for the PSI-analyzed 22-day-old heat-treated plants. **G.** RGB-based color-segmentation analysis showing distribution of pigmentation before and after heat stress for the 22-day-old heat-treated plants.

A

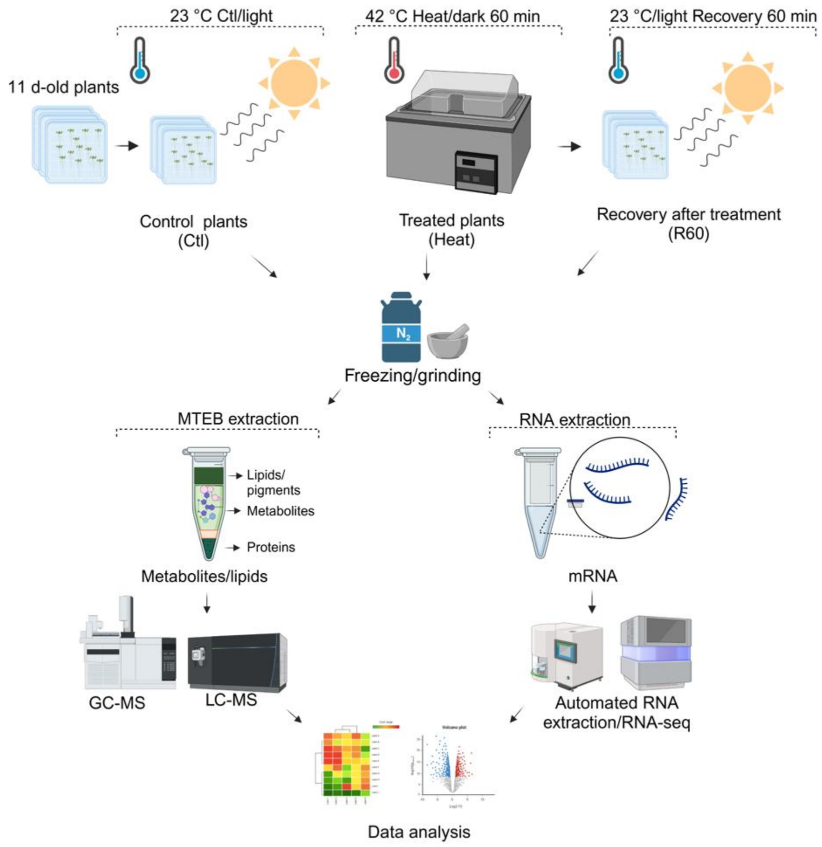


**Supplementary Figure 9. Design principle and experimental workflow of the multi-omics experiment**. Col-0 and *rbp47abcc'* plants were grown under control conditions (23°C, light) for 11 days. Plants were then subjected to heat stress (42°C, dark, 60 min) and subsequently allowed to recover under control conditions for 60 min. After treatment, samples were flash-frozen, ground in liquid nitrogen, and processed for multi-omics analysis. Metabolites, lipids, and proteins were extracted using the MTEB protocol, and their profiles were analyzed using GC-MS (metabolomics) and LC-MS (lipidomics). In parallel, mRNA was extracted, and transcriptomic profiling was performed using RNA sequencing (RNA-seq) on the Illumina platform. Data from all omics layers were integrated for further analysis.

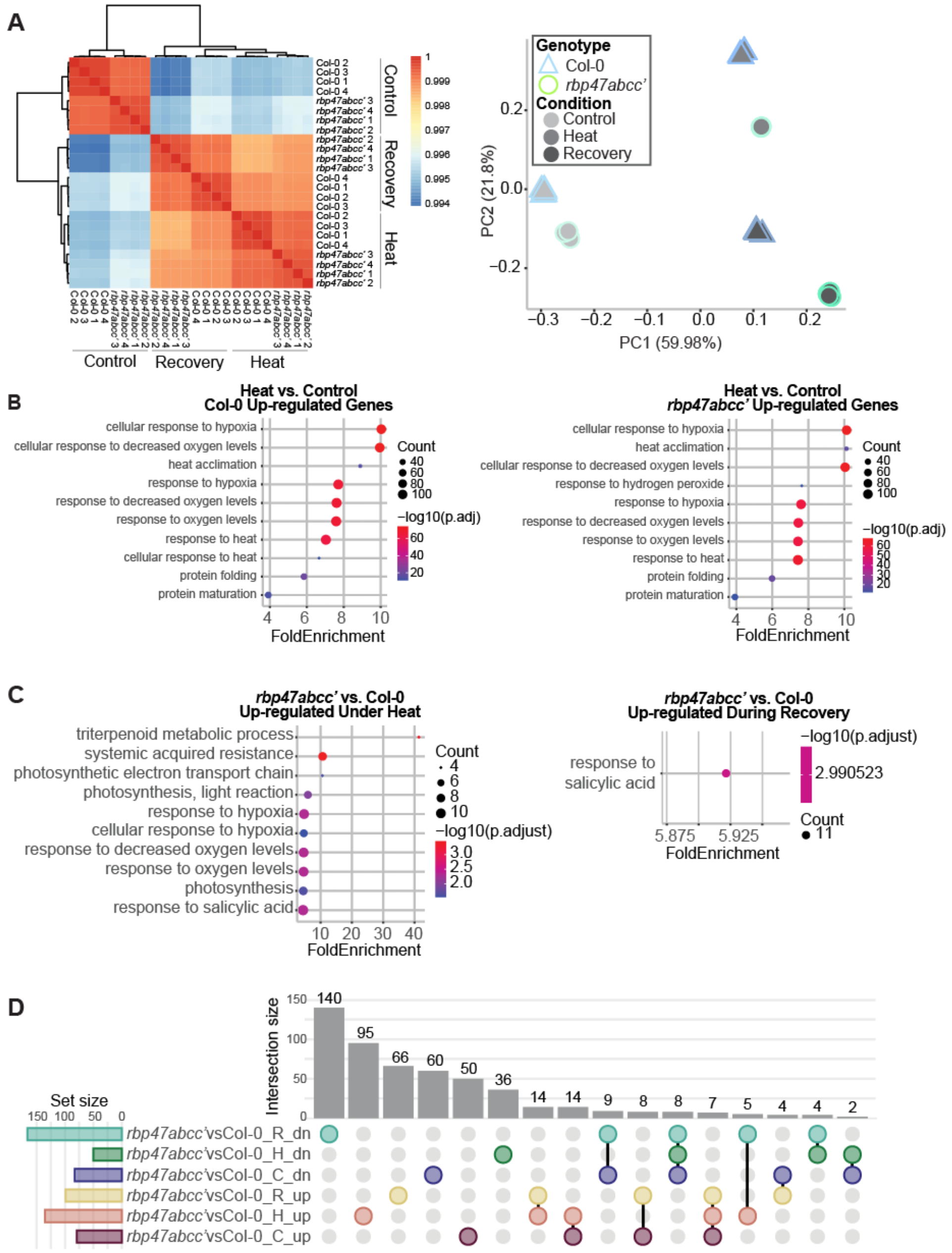


**Supplementary Figure 10. Characterization of the transcriptional response in Col-0 and *rbp47abcc'* under heat stress and recovery conditions. A.** Left: Hierarchical clustering of replicates indicate high reproducibility between samples. Right: Principal component analysis reveals a strong genotype (shapes) and treatment (colors) effect. **B.** GO enrichment of genes that are differentially abundant as a result of treatment (heat vs control) in either Col-0 (left) or *rbp47abcc'* (right). **C.** GO enrichment of genes that are up-regulated in *rbp47abcc'* compared to Col-0 in heat (left) or recovery (right) conditions. **D.** Upset plot describing the number of differentially abundant genes that are up or down in the mutant background relative to Col-0 across the different conditions.

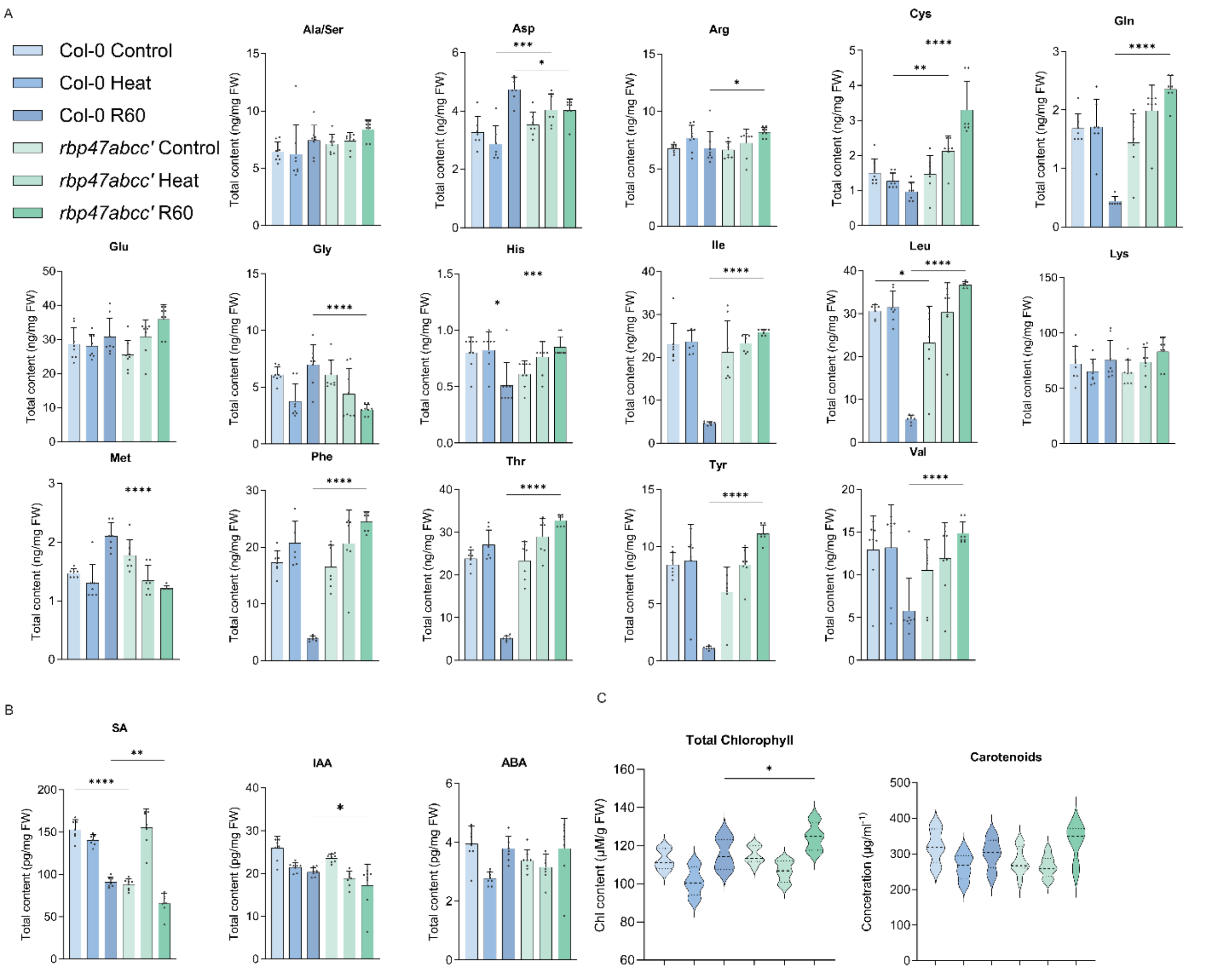


**Supplementary Figure 11. Target analysis of amino acid, hormones and pigments. A**. Amino acids were extracted from Col-0 and *rbp47abcc'* plants under control, heat stress (42°C, 60 min), and recovery (R60, 60 min after heat stress) conditions using the MTEB protocol. **B**. Quantification of the phytohormones Salicylic Acid (SA), Indole-3-Acetic Acid (IAA), and Abscisic Acid (ABA), extracted and analyzed as in **A**. **C**. Total Chlorophyll and Carotenoid content were measured under the same experimental conditions. Bar plots show mean ± SD ($n$ = 3, from three independent experiments). Statistical significance was assessed using one-way ANOVA followed by Dunnett's multiple comparisons test ($p < 0.05$; $p < 0.01$; *$p < 0.001$; **$p < 0.0001$).

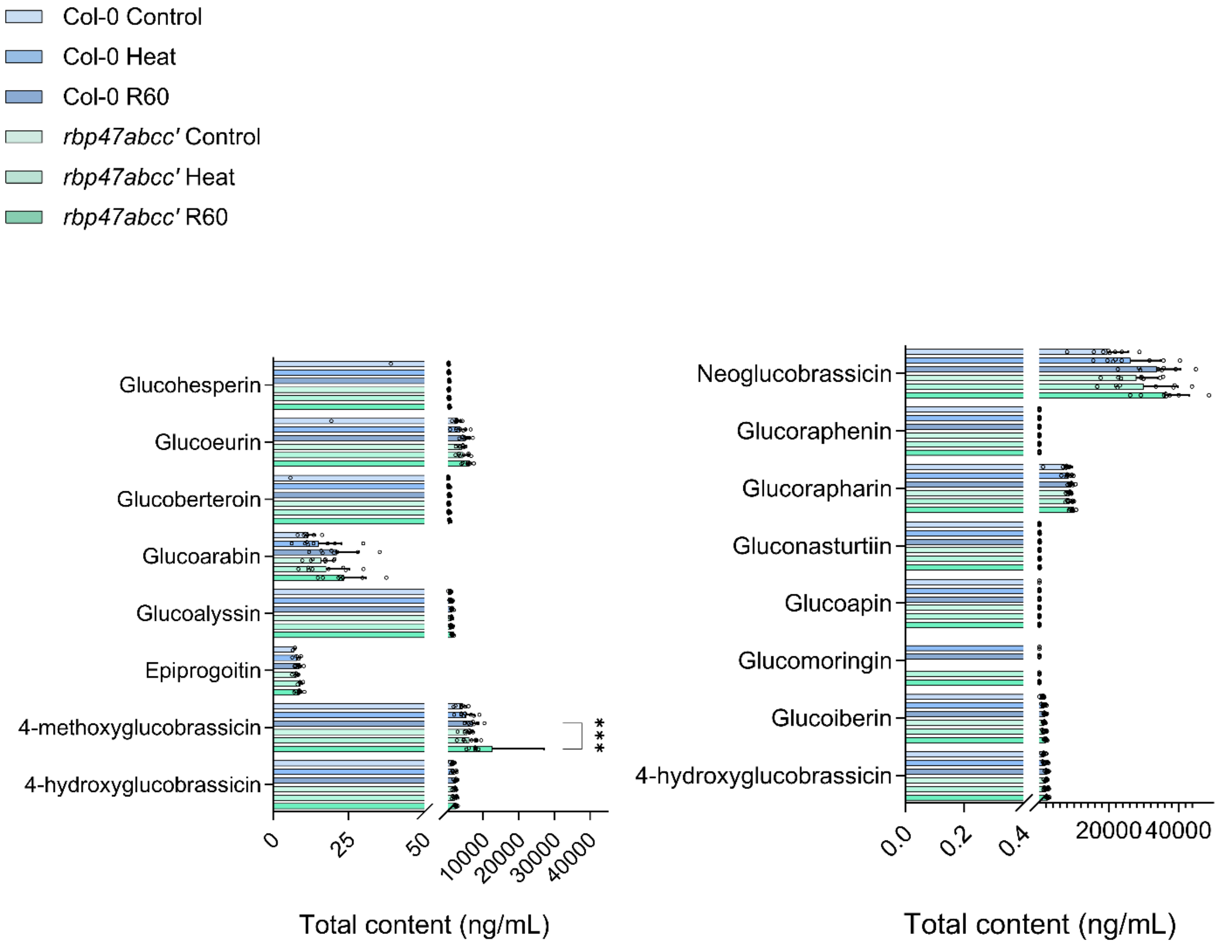


**Supplementary Figure 12. Target determination of Glucosinolates.** Glucosinolates were extracted from Col-0 and *rbp47abcc′* plants grown under control, heat stress (42°C, 60 min), and recovery (R60) conditions using the MTEB protocol. Their profiles were analyzed by GC-MS. Bar plots represent the mean ± SD ($n$ = 8, biological replicates). Statistical significance was determined using one-way ANOVA followed by Dunnett's multiple comparisons test ($p < 0.05$; $p < 0.01$; *$p < 0.001$; **$p < 0.0001$).
